# Dynamic microtubule-end structure governs multivalent kinetochore coupling by the Dam1c ring

**DOI:** 10.64898/2026.09.17.752406

**Authors:** Maksim Kalutskii, Helmut Grubmüller, Maxim Igaev

**Affiliations:** Theoretical and Computational Biophysics, Max Planck Institute for Multidisciplinary Sciences, 37077 Göttingen, Germany; School of Natural Sciences, Birkbeck, University of London, London WC1E 7HX, United Kingdom

## Abstract

Accurate chromosome segregation relies on kinetochores maintaining load-bearing attachments to dynamic microtubule ends, a coupling in which the Dam1 complex ring is central. How the microtubule-end structure and ring–microtubule interactions collectively determine attachment stability and drive force transduction remains unresolved. Here, we use multiscale modeling to show that the ring forms a “fuzzy” complex mediated by a dynamic network of intrinsically disordered regions, enabling both high-affinity binding and free diffusion along the microtubule lattice. Crucially, we find that this prototypic disordered–disordered protein complex provides most of the kinetochore–microtubule stabilization. By contrast, protofilament bending, commonly thought to drive force transduction, is sensitive to the arrangement of protofilaments along the ring and insufficient on its own to establish a robust coupling. The architecture of the dynamic microtubule end unifies these mechanisms by controlling both the progressive loss of fuzzy contacts and the resistance generated by protofilament bending. By accounting for the full conformational ensemble of microtubule structures, our model produces rupture forces similar to those previously measured and explains the distinct, tension-dependent behavior of kinetochore attachments to growing versus shortening microtubule ends. We propose that the Dam1 complex ring acts as a biased-diffusion coupler that probes and gradually remodels the evolving conformational landscape of the microtubule end under tension.

## Introduction

Faithful chromosome segregation relies on load-bearing attachments between centromeric kinetochores (KTs) and the ends of spindle microtubules (MTs).^1^ These attachments must be robust enough to withstand tension across bi-oriented chromosomes in metaphase, yet dynamic enough to remain coupled to shortening MT ends and translate their depolymerization into chromosome movement during anaphase.^2,3^ The budding yeast KT complex, which binds to a single MT end, is the best-characterized model system for studying this coupling.^2,3^ Physiological levels of tension on a KT attached to a growing MT end accelerate polymerization but weaken the attachment, while the same tension on a KT attached to a shortening MT end slows depolymerization and increases KT–MT attachment lifetimes – a phenomenon in mechanobiology usually referred to as a “catch bond”.^4–7^ This sensitivity and tension-dependent stabilization have been proposed to prevent accidental chromosome loss during segregation while facilitating error correction before erroneous attachments lead to chromosome abnormalities.^8,9^ Nevertheless, their molecular and physical origin remains unresolved.

The 10-protein Dam1 complex (Dam1c), which self-assembles into large rings around MTs,^10–15^ is a central component of the KT–MT interface in budding yeast and other fungi^16^ and a leading candidate for mediating this force-dependent attachment. The ring tracks shortening MT ends and couples MT depolymerization to movement under load, acting as a processive force coupler.^10,11,17^

How the ring turns MT dynamics into force is traditionally described by two classes of models. In power-stroke models,^18–21^ the outwards curling of protofilaments (PFs) pushes the non-diffusive ring towards the MT minus end. In biased-diffusion models,^17,22–24^ the ring freely diffuses along the MT lattice, and the flared shortening MT end poses a physical barrier that biases the ring motion. The underlying mechanisms are not necessarily mutually exclusive and may operate in combination. Both pictures assign a central role to the MT-end structure but make opposite assumptions about ring mobility and how the force is transmitted across the ring–MT interface.

Ring mobility is therefore a key observable for determining which mechanism dominates, yet reported estimates of the Dam1c ring’s diffusion coefficient span three orders of magnitude.^12,25,26^ Furthermore, single-molecule tracking of fluorescently labeled Dam1c oligomers on MTs shows that Dam1c diffusivity decreases sharply with oligomer size. While small oligomers are highly mobile,^25,27^ complete rings are nearly stationary, which would favor a power-stroke model. In contrast, cryo-electron microscopy (cryo-EM) and cryo-electron tomography (cryo-ET) analyses of ring conformations on MTs in vitro^24^ and in vivo^15^ point to low energy barriers for translational motion, hence supporting a biased-diffusion model. Whether the ring diffuses or is locked in place determines the force coupling mechanism, so resolving this discrepancy is a key to understanding how the Dam1c ring couples to a dynamic MT end.

Part of this discrepancy arises from how little is known about the KT–MT interface itself. Structural and cross-linking studies suggest that the interaction of the Dam1c ring with the MT is mediated primarily by its disordered regions.^28–34^ More specifically, these studies indicate that the Dam1 and Duo1 proteins of each Dam1c subunit contact the MT surface through long, disordered C-terminal tails (CTTs) spanning 193 and 72 residues, respectively.^28,29^ The CTTs of *α* - and *β* -tubulin, which provide interaction sites for numerous MT-associated proteins and post-translational modifications,^35^ have also been implicated, although their quantitative contribution remains controversial.^11,24^

How these CTTs engage with the MT lattice is difficult to determine by conventional structural methods. In particular, the conformations of intrinsically disordered regions are averaged out during cryo-EM density reconstruction, while cross-linking mass spectrometry (XL-MS) reports only residue pairs compatible with the cross-linker chemistry. As a result, these CTT-mediated interactions remain poorly understood, even though they are thought to contribute profoundly to the Dam1c mobility and affinity for the MT lattice.

Another unresolved question is whether CTT-mediated binding also contributes to the KT–MT force coupling. Tubulin CTTs can act as mechanical handles through which severing enzymes apply force to the MT lattice,^36,37^ demonstrating their capacity to bear mechanical loads. However, most models attribute the MT force on the Dam1c ring either to PFs releasing mechanical strain by curling outwards or to diffusion biased by the flaring MT tip, while the role of the CTT binding itself has received little attention. Yet, a ring attached to the MT lattice through many disordered regions must detach them progressively before it can leave the MT end, and this release could, in principle, also impede unbinding even if all PFs were straight. The complexity of this dynamic system is further increased by the observation that MT ends are flared and structurally heterogeneous,^38–41^ with individual PFs having unequal lengths and forming transient, laterally coupled clusters.^42–46^ Exactly how this heterogeneity in the MT-end structure translates into a mechanical barrier for the Dam1c ring also remains largely unexplored.

Here we use multiscale coarse-grained (CG) simulations to determine how the intrinsically disordered regions and the MT-end structure together produce and control the mechanical “grip” of the Dam1c ring. We first characterize the conformational dynamics of the ring–MT interface using a residue-level CALVADOS model^47–49^ and show that the CTTs of the Dam1 and Duo1 proteins in Dam1c form a dynamic electrostatic network with tubulin, particularly with its acidic CTTs. This multivalent interface features both high binding affinity and rapid contact turnover, allowing the ring to remain attached while freely diffusing along the MT lattice. Next, we quantify the resistance of the ring–MT interface to tension. By restraining all PFs in a straight conformation, we demonstrate that the CTT-mediated binding poses a substantial energetic barrier that confers unexpectedly high stability to the ring–MT interface even in the absence of PF curling. Further, we find that its magnitude depends on the MT-end structure: whereas blunt tips produce high resistance forces, extended or tapered tips provide a gradual path for sequential CTT unbinding and therefore produce lower resistance. Finally, using our recently published ultra-CG (UCG) model,^43^ we show that straightening bent and clustered PFs at the MT end is a second major stabilizing factor for the ring–MT coupling. Crucially, this mechanism requires that several long PFs be distributed evenly around the ring. In the absence of such an even distribution, the same long PFs grouped on one side of the ring, though a relatively rare configuration, instead provide a low-barrier escape route that facilitates ring detachment.

Together, these results show that the stability of the Dam1c ring–MT attachment under tension is an emergent property of the complex conformational ensemble and dynamics of the MT tip. Diffusion of the Dam1c ring mechanically probes the morphology of diverse tip configurations as the MT disassembles. Most of these configurations provide sufficient resistance, while ring detachment is dominated by rare but low-barrier geometries. Our results favor a biased-diffusion mechanism, in which tension applied to the Dam1c ring selectively stabilizes the attachment by remodeling the conformational landscape of the MT end.

## Results and discussion

### Structural insight into the Dam1c ring–MT interaction

To characterize the Dam1c ring–MT interface, we first constructed an atomistic model of the 16-heterodecamer Dam1c ring encircling the MT. The ring was assembled using the recently published coordinates of a Dam1c monomer (PDB ID: 8Q85^34^) and refined against the ring density extracted from the consensus reconstruction of the entire KT–MT complex using Correlation-Driven Molecular Dynamics (CDMD).^34,50^ The unresolved elements of this KT–MT complex included the CTTs of the Dam1 and Duo1 proteins, which span 193 and 72 residues, respectively, as well as the shorter CTTs of *α* - and *β* -tubulin.^33,34^ To include these unresolved regions explicitly, the atomistic ring–MT model was first converted to the CG representation, after which the Dam1, Duo1, and tubulin CTTs were added manually to their corresponding folded domains (Fig. 1a, left). The Ask1 CTT, another disordered domain of Dam1c, was not included because it does not bind the MT lattice and is outside the scope of this study. The CALVADOS package for implicit-solvent, residue-level CG modeling of biomolecules was used to simulate the dynamics of the ring–MT complex.^47–49^

**Figure 1:**
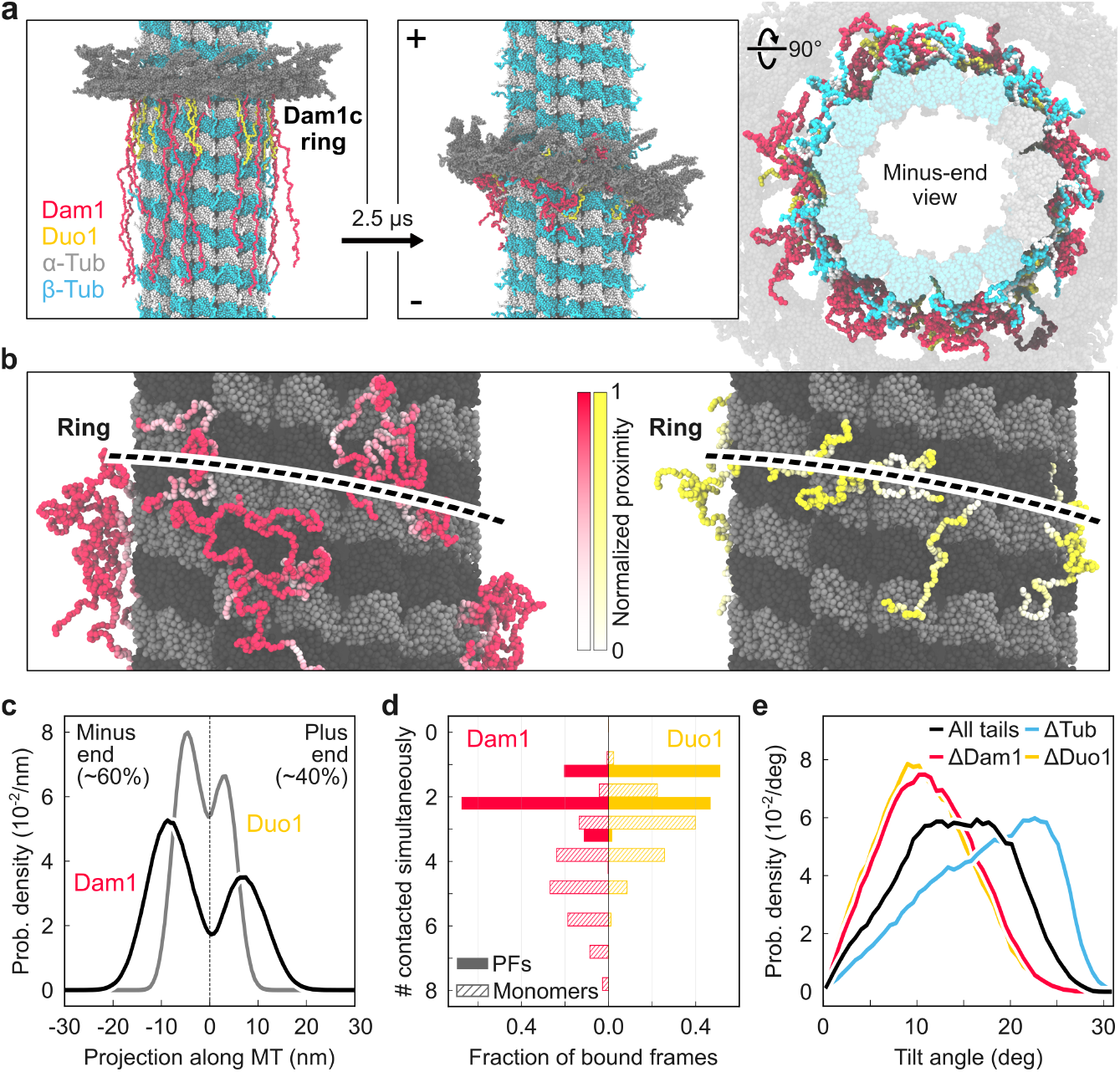
Architecture of the Dam1c ring–MT interface. **(a)** Initial configuration and a representative conformation after 2.5 *µ*s of a Dam1c ring encircling a MT (side and MT-end views). Dam1, Duo1, *α* -tubulin, and *β* -tubulin are shown in crimson, yellow, gray, and cyan, respectively. **(b)** Representative conformations of the Dam1 and Duo1 CTTs, with the residues colored according to their normalized proximity to the MT lattice. The dashed lines indicate the approximate position of the ring (hidden for clarity). **(c)** Probability densities of the projections of the Dam1 and Duo1 CTT end-to-end vectors along the MT axis and centered relative to the ring core. **(d)** Distribution of the numbers of PFs (solid) and tubulin monomers (hatched) simultaneously contacted by the Dam1 or Duo1 CTTs per frame. **(e)** Probability densities of the ring tilt angles for the wild-type complex (all CTTs present) and after removing the tubulin (ΔTub), Dam1 (ΔDam1), or Duo1 (ΔDuo1) CTTs.

During the initial equilibration simulation, the CTTs of the Dam1c ring, which were initially placed in extended conformations, rapidly condensed onto the MT surface beneath the ring (Fig. 1a, right; Fig. 1b). After equilibration, the CTTs remained highly dynamic and conformationally heterogeneous, with only a small fraction of their residues directly contacting the MT lattice in every simulation frame (Fig. 1b). Although all CTTs were initially oriented towards the MT minus end, individual tails repeatedly crossed the ring plane and reversed their axial orientation. At equilibrium, ∼60% of the Dam1/Duo1 CTTs pointed towards the minus end and ∼40% towards the plus end, indicating a weak directional preference (Fig. 1b; Fig. 1c). This dynamic interchange is consistent with cryo-EM observations showing no preferred orientation of the ring towards either MT end.^24^ We speculate that the modest 20% shift in orientational preference towards the MT minus end could facilitate the interaction between the Dam1 CTTs and the calponin-homology (CH)-domains of Ndc80 complexes (Ndc80c) aligned across the ring and pointing towards the MT minus end – another major component of the outer KT that has been shown to be essential for establishing robust KT–MT attachments and healthy chromosome segregation.^32,34^

In our simulations, the CTT–tubulin interactions were long-range and reached across several tubulin dimers. Projected along the main MT axis, the Dam1 and Duo1 CTTs extended by up to ∼20 nm and ∼10 nm, respectively (Fig. 1c). In addition, the two types of CTTs engaged differently with the MT lattice: each Dam1 CTT typically bridged 4–6 tubulin monomers on neighboring PFs, whereas each Duo1 CTT typically bound 2–4 tubulin monomers mostly within one PF (Fig. 1d).

Beyond the flexible CTTs, the Dam1c ring extends folded “bridge” domains from its core towards the MT surface. These bridges are formed by a coiled-coil motif of Dam1/Duo1 C-terminal helices and appear as prominent densities pointing towards the MT surface in cryo-EM studies.^13,24,30,33,34^ The slight bridge orientation towards the minus end may explain the modest orientational preference of the CTTs observed in our simulations.^24,34^ The folded bridges have also been proposed to be important in the power-stroke models.^18–20^ In our simulations, however, the folded bridges did not form any persistent contacts with the MT surface; instead, the disordered Dam1 and Duo1 CTTs formed a dense and continuously changing layer between the ring and the MT lattice (Fig. 1a, right; Movie S1).

Notably, the ring adopted a persistent tilt relative to the MT axis, with a mean angle of ∼14^°^ (Fig. 1e), consistent with early cryo-EM ensemble measurements.^13^ Our simulations indicate that this tilt is intrinsic to the geometry of the ring–MT interface: upon deletion of all CTTs, the ring still remained tilted by ∼12^°^ (Fig. S1). This residual tilt likely reflects the symmetry mismatch between the 16-heterodecamer ring and the 13-PF helical MT lattice, in which equivalent binding sites on neighboring PFs are axially shifted. The CTTs modulate this intrinsic tilt: removing the tubulin CTTs increased the mean tilt from ∼8^°^ to ∼12^°^ for the tail-free ring and from ∼14^°^ to ∼25^°^ when the Dam1 and Duo1 CTTs are present (Figs. 1e and S1). Thus, the flexible tubulin CTTs shield the ring from direct interactions with the helical MT lattice. The larger tilting effect in the presence of the Dam1 and Duo1 CTTs may reflect their asymmetric engagement with the MT interface, which reinforces the tilt arising from the ring–MT symmetry mismatch. Consistent with this picture, removing the Dam1 or Duo1 CTTs from the otherwise complete complex reduced the mean tilt from ∼14^°^ to ∼10^°^ and ∼8^°^, respectively (Fig. 1e). Thus, the mean non-zero tilt emerges from a dynamic balance between the ring–MT symmetry mismatch and the competing effects of the different CTTs.

Taken together, our results reveal a highly dynamic picture of the Dam1c ring–MT interface that deviates from the older view in which short ordered “bridges” serve as power-stroke couplers, similar to motor proteins, and from the way folded protein complexes typically assemble. Instead, the Dam1c ring engages with the MT surface through the disordered Dam1 and Duo1 CTTs, forming a dynamic layer with the tubulin CTTs.

### Tubulin C-terminal tails provide an electrostatic capture layer for Dam1c

Having established that the Dam1c ring engages with the MT predominantly through the disordered Dam1 and Duo1 CTTs, we next characterized the thermodynamic and kinetic properties of this interaction. The interacting CTTs have complementary charge distributions: the Dam1 and Duo1 CTTs are enriched in positively charged residues, whereas the tubulin CTTs are predominantly negatively charged (Fig. 2a). To determine how this electrostatic complementarity shapes the interface, we calculated residue–residue contact probability maps for all pairings between the Dam1/Duo1 CTTs and *α* -/*β* -tubulin. The resulting 2D maps revealed a multivalent interaction pattern characteristic of highly charged disordered regions (Fig. S2) and consistent with the dynamic interface observed in Fig. 1. Contact formation was governed by the interaction propensities of individual residues rather than specific residue-pair recognition (see Methods). To better illustrate the interaction patterns, we quantified the 1D marginal contact probabilities for each pair of CTT partners separately and identified regions contributing most to the Dam1c–tubulin interaction (see Fig. 2b for Dam1 and Fig. S3 for Duo1).

**Figure 2:**
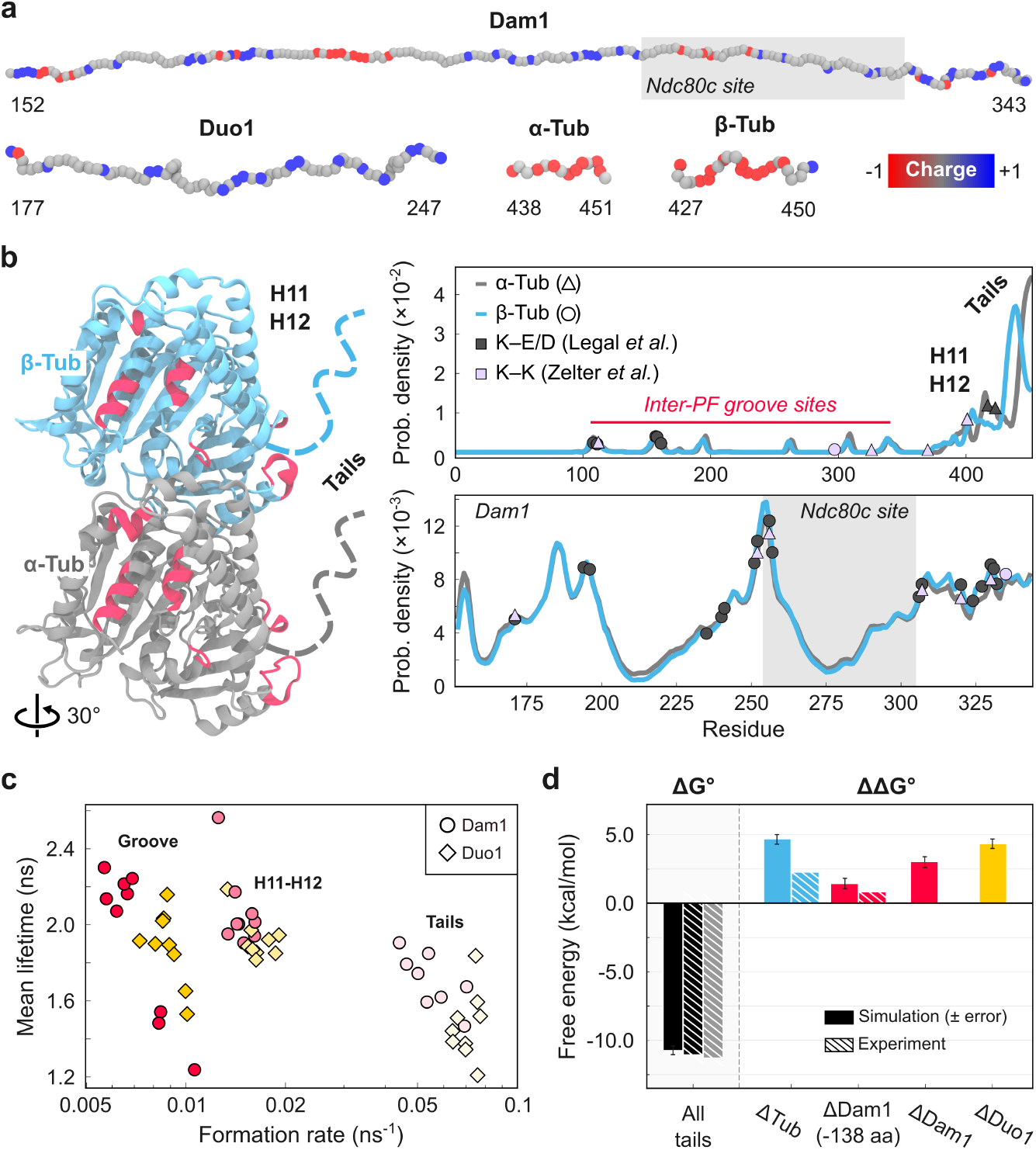
Thermodynamic and kinetic properties of the Dam1c–tubulin interaction. **(a)** Dam1, Duo1, and tubulin CTTs colored by residue charge. **(b)** Inter-PF groove interaction hotspots of the Dam1c CTTs mapped onto the *α* - (gray) and *β* -tubulin (cyan) structures and shown in crimson. Marginal contact probabilities are shown for tubulin (top) and the Dam1 CTT (bottom). Cross-linked residues identified by Legal *et al*. (K–E/D, dark gray)^29^ and Zelter *et al*. (K–K, light purple)^28^are marked by symbols. Shading indicates the Ndc80c-binding motif. **(c)** Kinetics of basic–acidic contacts with the tubulin groove, H11–H12 helices, and CTTs for Dam1 (circles) and Duo1 (squares). **(d)** Standard binding free energy Δ*G* ^0^ of wild-type Dam1c and changes ΔΔ*G* ^0^ upon CTT deletions. Solid-colored bars show simulation results, whereas hatched and gray bars show experimental estimates derived from previously reported binding affinities.^11,24,27^

The MT interactions can be grouped into three main regions (Fig. 2b). Approximately 16% of these contacts occurred within the inter-PF groove, ∼32% within the outer H11–H12 helices, and ∼52% within the tubulin CTTs. The acidic content increased in parallel across these regions, from ∼22% within the groove, to ∼35% within the H11–H12 helices, and ∼60% within the tubulin CTTs. As expected, the positively charged Dam1c CTTs most frequently interacted with acidic residues within the tubulin CTTs (*β* E438/D439/D440 and *α* E449/E450) and the H11–H12 helices (*α* E420/E423 and *β* E410/D417).

The identified interaction sites agreed well with the available cross-linking mass spectrometry data.^28,29^ EDC (carbodiimide) cross-links reported by Legal *et al*.^29^ mapped well to *α* E417/E423 in the H11–H12 helix region and *β* E157/E158/D161 in the inter-PF groove, as well as to the weaker *β* E108/E111 site. Similarly, lysine cross-links identified by Zelter *et al*.,^28^ located predominantly on the surface of *α* -tubulin, were very close to the contact peaks in our simulations. Notably, neither experimental dataset captured the dominant CTT interactions, possibly due to the poor detectability of long and highly negatively charged peptides with no basic residues in standard XL-MS workflows.

The Dam1 and Duo1 CTTs showed distinct, sequence-dependent interaction patterns. The Dam1 CTTs are organized into two contact-rich basic regions separated by an acidic insertion (Fig. 2a), which was strongly depleted from the anionic MT surface. The basic clusters located approximately within residues 151–200 and 231–343 corresponded to the peaks in the 1D contact probability profiles, whereas the acidic segment^207^ENEEDYEDD^215^ formed a pronounced contact frequency minimum. In contrast, the behavior of the more uniformly basic Duo1 CTTs resembled that of classical polyelectrolytes, with contact probabilities increasing towards their C-termini and no comparable acidic spacer (see Figs. 2b and S3).

The Dam1/Duo1 CTTs spent more than half of their time bound to the acidic tubulin CTTs. This high occupancy can arise either from long-lived contacts or from short-lived contacts that rapidly reform. To distinguish between the two scenarios, we computed the lifetimes and formation rates of representative basic–acidic contacts from the three interaction re (inter-PF groove, H11–H12 region, and tubulin CTTs; see Fig. 2c). Individual contacts were uniformly short-lived and produced single-exponential lifetime distributions. Mean contact lifetimes were similar in the inter-PF groove, H11–H12 region, and tubulin CTTs: ∼1.9, ∼2.0, and ∼1.6 ns, respectively, whereas mean formation rates increased by nearly an order of magnitude from the inter-PF groove to the tubulin CTTs (Fig. 2c). Thus, the tubulin CTTs dominate the Dam1c–tubulin interaction through frequent re-engagement rather than persistent contacts, creating an electrostatic layer that continuously rebinds the Dam1c CTTs. The nanosecond interaction timescale is consistent with other high-affinity complexes formed by oppositely charged disordered regions, such as ProT*α* –H1.^51,52^

We next characterized the thermodynamics of the Dam1c–tubulin binding. We calculated the standard binding free energy of a single Dam1c subunit binding to the MT while selectively removing individual CTTs from the binding interface (Fig. 2d). The wild-type complex bound the MT lattice with Δ*G* ^0^ ≈ −11 kcal mol^−1^ (∼18.5 *k*_B_*T*), consistent with nanomolar affinities reported previously.^10,11^ Removal of the tubulin CTTs strongly reduced the affinity (ΔΔ*G* ^0^ ≈ 4.7 kcal mol^−1^), in agreement with their dominant contribution to the interaction network. Removal of 138 proximal Dam1 CTT residues only moderately weakened the binding (ΔΔ*G* ^0^ ≈ 1.4 kcal mol^−1^), which was consistent with observations that partially truncated Dam1 CTT constructs largely retain MT affinity.^24,53^ Removing the full 193-residue Dam1 CTTs weakened MT binding by approximately two-thirds as much as removing the tubulin CTTs (ΔΔ*G* ^0^ ≈ 3 kcal mol^−1^). Removing the Duo1 CTTs produced a strong weakening effect comparable to that of removing the tubulin CTTs (ΔΔ*G* ^0^ ≈ 4.3 kcal mol^−1^). We attribute this strong contribution to the shorter length and more uniformly basic charge distribution of the Duo1 CTTs (Figs. 1b and 2a), which may allow favorable electrostatic interactions with a smaller loss of conformational entropy upon binding. This finding further supports the hypothesis that Duo1 can mediate MT binding when the Dam1 CTTs are removed^24^ or inhibited, *e*.*g*., by Aurora B phosphorylation.^2,3^

Together, these values provide a quantitative decomposition of the Dam1c binding affinity for the MT surface. More broadly, our results place the Dam1c–MT interface within the emerging class of high-affinity, dynamic complexes formed by oppositely charged intrinsically disordered regions.^51,52^

### The Dam1c ring freely diffuses along the microtubule surface

The mobility of the Dam1c ring on the MT lattice determines how the complex couples to dynamic MT ends, but the physical basis of this mobility remains unresolved. Diffusion coefficients of the ring measured in vitro span three orders of magnitude.^12,25,26^ Cryo-EM analyses of the ring tilt and spacing imply a translational barrier of approximately 1 *k*_B_*T* per diffusion step, consistent with a freely diffusing ring,^15,24^ whereas Dam1c-oligomer diffusion decreases rapidly with size and extrapolates to an essentially immobile ring, supporting a diffusion-free forced walk.^26^ The two views agree in that the ring binds tightly, but differ in whether this tight binding implies immobility (Fig. 3a).

**Figure 3:**
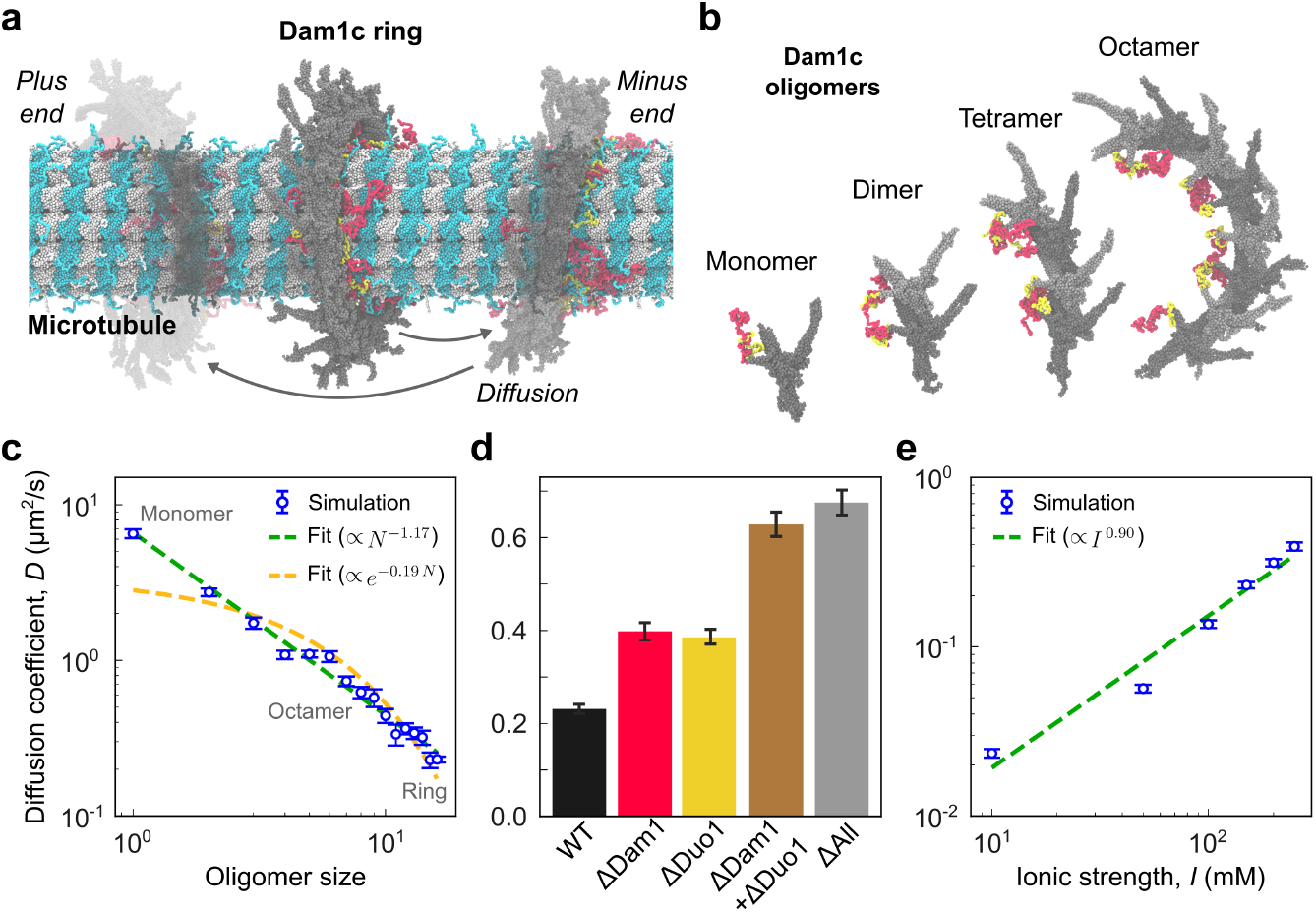
Dam1c oligomers freely diffuse along the MT lattice. **(a)** Simulation snapshot of the Dam1c ring on the MT (plus and minus ends indicated). Arrows denote 1D lattice diffusion. **(b)** Simulation snapshots of Dam1c oligomers of increasing size (monomer, dimer, tetramer, and octamer). **(c)** Diffusion coefficient *D* of Dam1c oligomers as a function of size *N*, from a monomer to a complete ring, with a power-law fit (∝ *N* ^−1.17^, green dashed line) and an exponential fit (∝ *e*^−0.19*N*^ , orange dashed line). **(d)** Diffusion coefficient *D* of the wild-type Dam1c ring and after removing the tubulin CTTs (ΔTub), Dam1 CTTs (ΔDam1), Duo1 CTTs (ΔDuo1), both basic CTTs (ΔDam1 + ΔDuo1), or all CTTs (ΔAll). **(e)** Diffusion coefficient *D* of the Dam1c ring as a function of ionic strength *I* (simulation, blue points with error bars; power-law fit *D* ∝ *I*^0.90^, green dashed line).

Because our model accurately reproduces the high affinity of a Dam1c monomer for the MT lattice (Fig. 2d), we next asked whether such tight binding is compatible with diffusion of the assembled ring (Fig. 3b). In our simulations, the ring remained diffusive with *D* ≈ 0.2 *µ*m^2^ *s* ^−1^, while a single Dam1c monomer diffused at ∼6.5 *µ*m^2^ *s* ^−1^. Both values are about an order of magnitude larger than the upper bound of the corresponding experimental estimates. We attribute this systematic offset to the smoother energy landscape of the CALVADOS force field.^47–49^ Although we rescaled time by benchmarking the translational diffusion of single-chain intrinsically disordered proteins (see Methods), this correction factor is system-dependent and has been observed to vary by up to three orders of magnitude.^54^ The absolute diffusion coefficients therefore carry substantial uncertainty.

To gain further insight into the physics of ring diffusion, regardless of the absolute timescale, we focused on how *D* scales with oligomer size. We computed the 1D diffusion coefficient *D* of Dam1c oligomers of increasing size (Fig. 3b,c) from the mean squared displacement of their center of mass along the MT axis (see Methods). *D* decreased as a power law of oligomer size *N, D* ∝ *N* ^−1.17^ (*R*^2^ = 0.98), whereas a single-exponential function did not fit the same data as well, *D* ∝ *e*^−0.19*N*^ (*R*^2^ = 0.47), and underestimated the mobility of small oligomers. The power-law dependence with an exponent close to −1.0 suggested additive per-subunit friction with little cooperativity between subunits. In this case, addition of a monomer would contribute a constant drag *ζ*, so that the total friction *ζ*_tot_ = *N ζ* and *D* ∝ *N* ^−1^. This is consistent with the “fuzzy” interface,^55^ in which the disordered CTTs form a highly redundant network of short-lived contacts (Fig. 2c).

To identify which parts of the interface cause drag, we computed *D* of the wild-type Dam1c ring versus modified rings lacking specific CTTs (Fig. 3d). Deleting either the Dam1 or the Duo1 CTTs roughly doubled *D* (to ∼0.4 *µ*m^2^ *s* ^−1^ in both cases), even though the Dam1 CTTs are more than twice as long as the Duo1 CTTs. This suggests that the drag is determined by the CTTs’ charge rather than their length, as the two types of CTTs carry an identical net charge of +15.

Consistently, deleting both basic CTTs together increased *D* to ∼0.65 *µ*m^2^ *s* ^−1^, essentially to the value obtained when all CTTs including the tubulin ones were removed (∼0.7 *µ*m^2^ *s* ^−1^). When both Dam1 and Duo1 CTTs were deleted, the tubulin CTTs contributed almost no further drag: the acidic tubulin CTTs slow the ring’s diffusion only through their contacts with the basic Dam1 and Duo1 CTTs, which are the physical carriers of the friction. Consistent with this electrostatic picture of the drag, *D* increased with ionic strength as *D* ∝ *I*^0.90^ (Fig. 3e), because stronger electrostatic screening reduces the drag on Dam1c oligomers.

Our simulations yield a diffusion coefficient of the Dam1c ring that falls into a fast diffusion regime. The ring is therefore quite mobile and remains so despite the high MT binding affinity (Δ*G* ^0^ ≈ −11 kcal mol^−1^ per Dam1c monomer). The tight binding and rapid diffusion coexist because the interface is multivalent and its contacts are transient, which we propose here to be the molecular basis for the decoupling of binding from mobility. Such behavior is compatible with a biased-diffusion mechanism, in which the depolymerizing MT end biases an otherwise unbiased random walk.

### Microtubule-end architecture determines the mechanical resistance of the Dam1c ring to unbinding

We next asked how unbinding of the Dam1c ring from a dynamic MT end is prevented. Two structurally distinct mechanisms are conceivable. First, the disordered Dam1/Duo1 CTTs that mediate MT binding could progressively detach as the ring leaves the lattice. In this case, it would be the multivalent interface characterized above that creates an energy barrier to ring unbinding. Second, PFs curling outwards at a flared MT end could sterically oppose the ring’s motion towards the MT plus end. Here, we refer to these two mechanisms as the CTT- and curvature-mediated resistance, respectively.

To estimate the strength of these contributions, we calculated potentials of mean force (PMFs) of our ring–MT system as a function of displacement of the Dam1c ring along the MT axis (Fig. 4a). For each profile, we then estimated the maximum mean resistance force ⟨*F*⟩_max_ as the steepest slope of the PMF. This quantity describes the load at which the barrier against ring detachment vanishes. We asked how this load depends on MT-tip architecture, which we described by the number of tubulin dimers by which each PF extends beyond the last complete lattice layer (Fig. S4a). Because the space of possible tip structures is combinatorially large, we separated individual structural degrees of freedom and analyzed their contributions independently.

**Figure 4:**
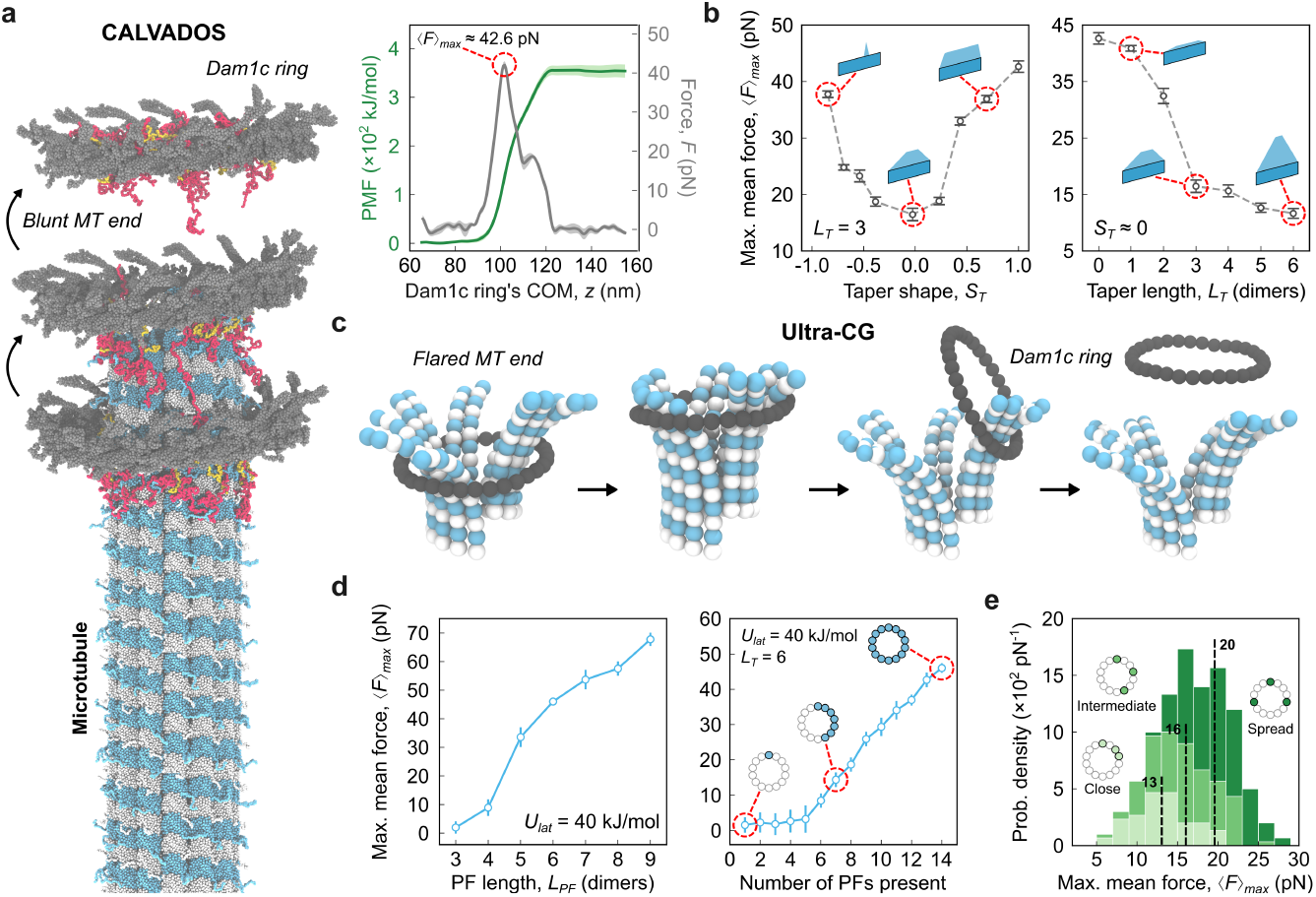
MT-end architecture controls the Dam1c ring’s unbinding barrier. **(a)** CTT-mediated unbinding from a straight MT end (CALVADOS model). Left, simulation snapshot of the Dam1c ring leaving a blunt MT end. Right, the corresponding PMF (green line with shaded areas reflecting statistical uncertainty) and the mean resistance force ⟨*F*⟩ (gray line with shaded areas reflecting statistical uncertainty) versus the ring’s center-of-mass position *z* . The peak force is ⟨*F*⟩_max_ ≈ 43 pN. **(b)** CTT-mediated ⟨*F*⟩ _max_ versus the MT-tip taperness *S*_*T*_ at a fixed taper length *L*_*T*_ = 3 dimers (left) and versus the taper length *L*_*T*_ for a fixed taperness *S*_*T*_ ≈ 0 (right). Cartoons illustrate unfolded 2D representations of MT lattices with the corresponding tip profiles. **(c)** Curvature-mediated unbinding from a flared MT end (UCG model). PFs at the MT end curl against the encircling Dam1c ring and partially straighten upon ring unbinding. **(d)** Curvature-mediated ⟨*F*⟩ _max_ for GDP-bound MT ends (*U*_lat_ = 40 kJ/mol) as a function of PF length *L*_PF_ with all PFs having an equal length (left), and as a function of the number of long PFs present (*L*_PF_ = 6 dimers; right). Cartoons illustrate the arrangement of long PFs along the MT circumference. **(e)** Distributions of curvature-mediated ⟨*F*⟩ _max_ over 300 randomly generated GDP-MT-end configurations with a fixed taperness *S*_*T*_ and colored by spatial organization of their longest PFs (“close”, “intermediate”, “spread”). Dashed lines mark the subpopulation means (∼13, ∼16, ∼20 pN, respectively).

We first investigated the CTT-mediated resistance using the residue-level CALVADOS model with all PFs restrained in straight conformations, thereby excluding any mechanical contribution from PF bending and focusing only on the resistance generated by progressive release of the multivalent Dam1c–MT interface. To test whether the arrangement of PFs shapes this escape barrier, we permuted PFs while keeping their lengths fixed. The observation that ⟨*F*⟩_max_ changed by only 1–3 pN (Fig. S5) indicates that CTT-mediated resistance is largely insensitive to how the PFs are arranged along the MT.

Next, we asked how the distribution of PF lengths affects this resistance. To quantify changes in MT-end shape, we introduced a shape parameter, *S*_*T*_ (see Methods). Briefly, *S*_*T*_ describes the distribution of PF lengths relative to the longest PF at the MT end. Near-zero *S*_*T*_ corresponds to PF lengths distributed rather evenly between the base and the maximum height. Positive *S*_*T*_ indicates that most PFs extend close to the maximum height, and negative *S*_*T*_ indicates that most PFs terminate close to the base (see schematics in Fig. 4b, left). At a fixed maximum PF length (taper length), *L*_T_, CTT-mediated ⟨*F*⟩ _max_ showed a pronounced U-shaped dependence on *S*_*T*_ (Fig. 4b). The force was lowest for an evenly tapered MT end (⟨*F*⟩ _max_ ≈ 16 pN) and increased symmetrically towards both extremes, reaching ⟨*F*⟩ _max_ ≈ 43 pN at a fully blunt tip. This finding indicates that the mechanical resistance of the ring is mainly governed by how gradually the CTTs unbind at the MT tip. Consistent with this picture, increasing the taper length *L*_*T*_ strongly extended the range over which the CTTs would need to unbind and reduced ⟨*F*⟩ _max_ (Fig. 4b, right). Most of this decrease occurred within the first three tubulin layers, where ⟨*F*⟩ _max_ dropped from ∼42 pN to ∼16 pN. Further increasing the taper length produced only a modest decrease, reaching ∼13 pN at six tubulin layers.

We then investigated the curvature-mediated contribution using our previously published UCG model of MT-end dynamics^43^ (Fig. 4c; see Methods). We used the UCG model because elastic networks combined with residue-specific CG models such as CALVADOS do not accurately capture large conformational changes of folded proteins,^56^ whereas our UCG model was specifically parameterized to reproduce PF dynamics. Curvature-mediated ⟨*F*⟩_max_ increased steeply with *L*_PF_, which was kept equal for all PFs (Fig. 4d, left). MT ends with *L*_PF_ ≤ 3 dimers produced almost no resistance (⟨*F*⟩ _max_ ≤ 3 pN), suggesting that only MT ends with *L*_PF_ ≥ 4 dimers are able to restrict the ring’s motion. For *L*_PF_ ≥ 5 dimers, the force became substantial (>30 pN), reaching ∼70 pN at *L*_PF_ = 9 dimers. These results show that long PFs are required to generate large resistance when they engage with the ring symmetrically.

Long PFs grouped on one side interact with the ring asymmetrically, which can tilt it rather than lock it in. We therefore asked how the force changes as the number of long PFs engaging the ring is reduced. At least six long PFs grouped together were required to produce a substantial resistance force of ∼10 pN or higher, while ⟨*F*⟩ _max_ gradually increased to ∼50 pN with more long PFs grouped at the MT end (Fig. 4d, right). Together, these results indicate that curvature-mediated forces are sensitive to both PF length and arrangement.

Given the sensitivity to the MT-tip morphology, we next asked what resistance is generated by an average disassembling MT end. To obtain a realistic ensemble of MT-tip compositions, we generated 10^5^ tip structures by sampling PF lengths from the distribution measured in our previous cryo-ET study.^43^ The resulting tip configurations were evenly tapered with *S*_*T*_ = −0.11 ± 0.25 (see Fig. S6d) and contained relatively few long PFs on average: ∼3.3 PFs of *L*_PF_ ≥ 5 dimers and ∼1.8 PFs of *L*_PF_ ≥ 6 dimers (Fig. S6b,c). We selected a representative tip composition containing three long PFs of 8, 6, and 5 dimers (*S*_*T*_ = −0.25) and calculated ⟨*F*⟩ _max_ for 300 random PF arrangements (Fig. 4e). Despite the identical set of PF lengths, the forces varied from ∼5 to ∼28 pN, with a mean of ∼17.0 pN. One-third of the arrangements produced force values of less than 15 pN. This broad distribution reflects the dominant contribution of the few long PFs to the resistance force. Across 300 arrangements, the force ⟨*F*⟩ _max_ increased with the spread of the long PFs (Pearson correlation of *r* ≈ 0.58), while the PMF height showed an even stronger dependence on their spatial distribution (*r* ≈ 0.87). The lowest forces were obtained when the long PFs were grouped on one side of the ring. Furthermore, placing PFs of similar length closer together reduced ⟨*F*⟩ _max_, whereas doing the opposite increased it (*r* ≈ 0.54). Typical MT ends therefore generate a broad range of resistance forces, determined by how their few long PFs are distributed around the ring.

Because 76% of tips contained only 1–4 long PFs (Fig. S6), we expect a similar distribution of resistance forces to hold across the full ensemble of possible tip configurations. We therefore consider this distribution to be characteristic of the curvature-mediated resistance generated by disassembling MT ends.

The maximum mean forces computed above correspond to barrier-vanishing loads and should be treated as upper bounds on the resistance forces measured experimentally.^4,5,32,34^ Under linearly increasing loads applied over experimentally relevant timescales, the ring escapes the MT tip by thermal activation once the barrier is sufficiently lowered. Combining the CTT- and curvature-mediated profiles for flared ends (see Methods) and using a Kramers-type model,^57,58^ *k*_off_ (*F*) = *k*_att_ exp[−Δ*G* ^‡^ (*F*)/*k*_B_*T* ], we define the rupture force as the constant load at which 1/*k*_off_ = 1 s as *F*_1*s*_ . Across the 300 tip arrangements, the rupture forces span a broad range, with a mean of ∼24 pN and maximum values of ∼33 pN (Fig. S7), comparable to the largest forces reported in optical-trap assays.^5,26,32,34^ Because a diffusing Dam1c ring is expected to sample multiple MT-end configurations before detachment, the experimentally relevant quantity is not the mean of the individual rupture forces, but the off-rate averaged over the ensemble of MT-end configurations, *k*_att_⟨exp(−Δ*G* ^‡^/*k*_B_*T*)⟩. This average is weighted towards low-barrier configurations and predicts an ensemble mean rupture force of ∼15 pN, in good agreement with the average experimental value of ∼10 pN.^5,26,32,34^

Together, these results suggest that the mechanical stability of the ring–MT attachment emerges from an interplay between the multivalent CTT interface and PF mechanics, and that the resistance to ring unbinding is controlled by the ensemble of MT-end structures.

## Conclusions

Using multiscale CG simulations, we have characterized how the Dam1c ring interacts with the MT lattice and how its attachment to a dynamic MT end can withstand tension and detachment. This interaction is primarily mediated by the disordered Dam1/Duo1 and tubulin CTTs, which together form a multivalent, electrostatically driven interface. This dynamic interface allows the ring to combine two seemingly contradictory functional properties: to diffuse rapidly along the MT lattice while remaining strongly bound to the MT surface.

The main result of our analysis is that the force sustained by this attachment strongly depends on MT-tip geometry. Both the CTT- and the curvature-mediated contribution to this resistance vary with the length distribution and spatial arrangement of PFs at the MT end. The two contributions to this grip respond differently to changes in the MT-end structure. The CTT-mediated resistance forces are determined by how gradually Dam1c–tubulin contacts are lost as the ring leaves the MT end. Blunt ends impose an abrupt termination of the binding surface and produce the largest resistance forces, whereas extended or tapered tips provide a more gradual path for contact release and therefore substantially reduce the force. Importantly, this interaction alone confers sufficient resistance to ring unbinding even in the absence of PF curling.

PF curling provides a second, mechanically distinct contribution. Long and curled PFs impose a mechanical barrier to ring unbinding, but only when several long PFs are distributed across the full perimeter of the ring. In contrast, when the same long PFs are grouped on one side, the ring can tilt and bypass them, thereby strongly reducing the barrier. Together, these results establish Dam1c attachment stability as an ensemble property of heterogeneous MT-end architectures, with the off-rate dominated by rare but low-barrier configurations. Across an ensemble of PF arrangements for an average disassembling MT tip, we obtain a maximum rupture force of ∼33 pN and an ensemble average of ∼15 pN, which agrees well with optical-trap experiments.^5,26,32,34^ The somewhat higher predicted forces may partly result from combining the CTT- and curvature-mediated contributions under the simplifying assumption that they are additive. A unified CG model that combines a residue-level description of CTT interactions with an accurate representation of PF mechanics would capture coupling between these contributions directly and could improve quantitative agreement with experimental rupture forces.

The sensitivity of curvature-mediated forces to MT-end geometry indicates that PF curling alone is insufficient to maintain robust KT–MT attachment. This points to a more general principle of KT–MT coupling: stable attachment emerges from rapidly exchanging, multivalent interactions that can adapt to an evolving MT tip. Like yeast Dam1c, human Ndc80 and Ska complexes also form multivalent assemblies at dynamic MT ends that span multiple tubulin subunits and neighboring PFs.^59–62^ Despite their evolutionary and structural differences,^16^ these couplers may therefore stabilize KT–MT attachments through the same physical principle.

This geometry-dependent picture also makes rapid Dam1c diffusion a key component of the coupling mechanism. During MT depolymerization, the ring can diffuse and rapidly re-equilibrate with newly formed MT-end configurations, allowing it to sample multiple MT-end architectures before detachment. Our results consistently favor a biased-diffusion mechanism of force coupling.^22,23^ The Dam1c ring behaves as a Brownian coupler whose motion is constrained by the depolymerizing MT end. Our simulations show neither persistent Dam1c–tubulin contacts nor strongly suppressed ring diffusion, arguing against a power-stroke mechanism. Instead, ring diffusion is rectified by curved PFs and by the gradual loss of individually weak CTT contacts, the latter being consistent with the Hill’s sleeve model.^22^ However, because our analysis is based on equilibrium free-energy landscapes rather than on the kinetics of MT depolymerization, we cannot fully rule out possible power-stroke contributions.

The geometry-dependent attachment seen in our simulations also explains the apparently stronger binding of the Dam1c ring to growing, mostly GTP-bound MT ends.^5^ Our previous structural work^43^ demonstrated that growing MT ends feature shorter and more clustered PFs, which more closely resemble a blunt tip as compared with shortening MT ends featuring longer but more independent PFs. Here we argue that these very distinct structural features can explain the different ring–MT attachment stability depending on the polymerization state of the MT end.^5^ The longer lifetimes of Dam1c attachments to growing MT ends therefore need not imply a higher affinity for GTP-tubulin, as tubulin conformation is not uniquely determined by nucleotide state but can be easily overridden by the cellular environment.^63^ Instead, the nucleotide state can regulate the Dam1c ring coupling indirectly by affecting the characteristic architecture of the MT end.

The same principle provides a statistical-mechanical explanation for the stabilization of Dam1c ring–MT attachments under tension. Our results suggest that tension applied to the ring shifts the ensemble of MT-tip configurations sampled during depolymerization. By suppressing PF curling, increasing tension biases the tip towards shorter and blunter configurations, thereby strengthening the CTT-mediated contribution to attachment stability (Fig. 5). This would reduce the probability of weak-grip configurations and increase the mean attachment lifetime. Such catch-bond-like behavior^4–6^ could therefore emerge from tension-dependent remodeling of the conformational ensemble of the MT end, independently of Aurora B phosphorylation. Additional support for MT-tip remodeling comes from the tension stabilization of MT disassembly.^5,26,64^ Consistent with this picture, suppression of MT disassembly by the addition of Ndc80c and Ska complexes was shown to remodel shortening MT ends in the proposed direction, namely, by producing shorter and more clustered PFs.^59^

**Figure 5:**
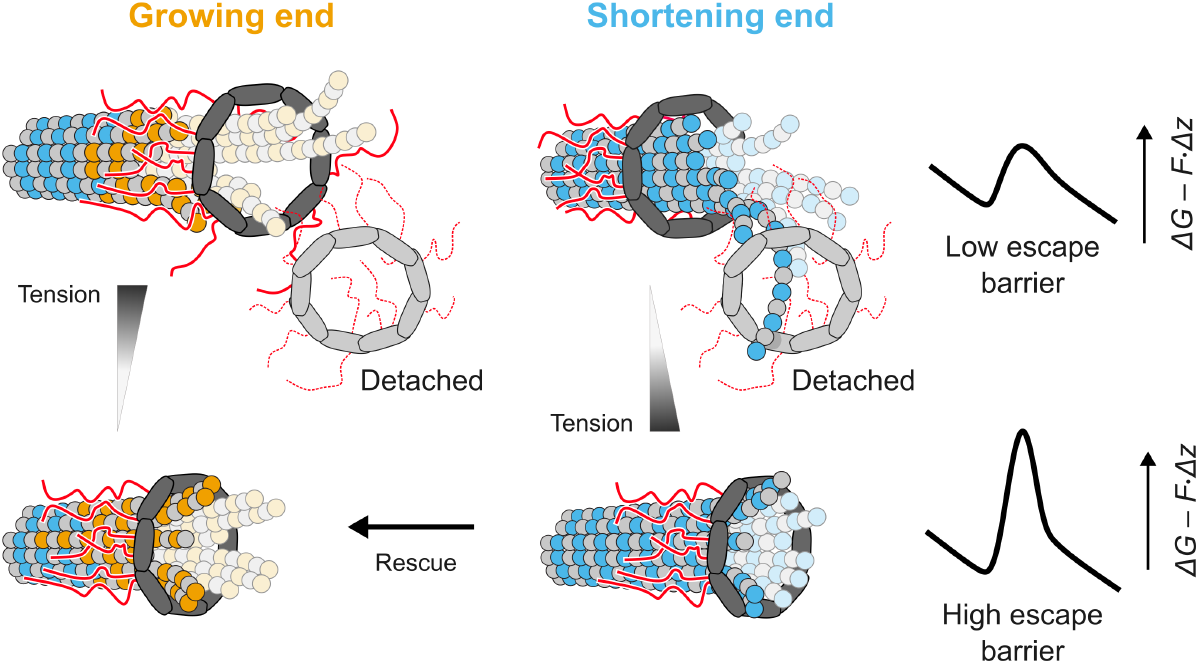
Schematic illustration of the force-induced changes of MT tip. Dam1c remains coupled to dynamic MT ends through a multivalent CTT-mediated interface, while the instantaneous tip architecture sets the barrier to ring escape. Tension stabilizes shortening ends by making them blunter and destabilizes growing ends by promoting more ragged tip structures.

Likewise, tension may remodel a growing MT end in the opposite manner. Faster MT assembly is typically associated with increased tip raggedness.^39,65^ The ability of the Dam1c ring to accelerate MT assembly under tension^4^ suggests that it may promote a more ragged architecture with distorted distributions of PF lengths and arrangements (Fig. 5). Such remodeling would lower the energy barrier for ring unbinding and could contribute to the slip-bond-like behavior of KT–MT attachments observed at growing ends.^5^ Thus, the opposite tension responses of Dam1c-ring attachment lifetimes to growing and shortening MT ends may arise from opposite tension-induced changes in the MT-end architecture.

Although we attribute MT-tip remodeling and the resulting tension-dependent stabilization of the KT–MT complex to the Dam1c ring, Dam1c is not directly coupled to the centromere. Force transmission between the outer and inner KT therefore relies on linker complexes such as Ndc80c. Fully understanding KT–MT coupling will thus require unraveling the complex interplay among Ndc80c, Dam1c, and the MT end. One prominent example of the consequences of this interplay is the effect of Stu2, which is required for the catch-bond-like behavior of KT–MT attachments.^64^ Miller *et al*. showed that Stu2 strengthens KT–MT attachments and proposed that this effect stems from enhanced Ndc80c–MT binding. We support this hypothesis and speculate that Stu2 enables, but does not generate, tension-dependent stabilization by supporting force transmission between the Dam1c ring and the centromere. Without Stu2, Ndc80c couples weakly to the MT surface^64^ and may therefore become the weak link, unbinding before force is efficiently transmitted from the Dam1c ring. This would explain both the prominent slip-bond behavior of Stu2-depleted attachments at growing and shortening MTs and the increased stability of these attachments under low force at shortening MTs. The latter may arise because weaker Ndc80c–MT coupling increases Ndc80c mobility, facilitating its redistribution and tracking of the disassembling MT end.

Taken together, our results identify the composition of the MT end as a key functional determinant that fine-tunes its multivalent attachment to the Dam1c ring. The dynamic CTT-mediated interface enables both strong attachment and rapid diffusion, which allows the ring to mechanically probe and adapt to the ever-changing ensemble of MT-end configurations. Its instantaneous architecture determines the energy barrier to ring unbinding and therefore the force that can be transmitted across the Dam1c ring–MT interface.

## Methods

### Construction of initial models

To construct the initial 13-PF GDP-bound MT models, we followed our previously published protocol for CDMD refinement.^42,50^ Briefly, initial coordinates for the tubulin dimers were obtained from PDB ID: 7SJ7. The missing loop in the *α* -subunit (residues 38 to 46; excluding the His-tag) and unresolved parts of the flexible C termini (*α* :442–451 and *β* :440–450) were modeled in for structure completeness using Modeller version 10.6^66^ but excluded from further refinement. The 3.8-Åreconstruction of an undecorated 13-PF wild-type GDP-MT from recombinant human tubulin (EMD-25156)^67^ was used to construct the all-atom (AA) model of a MT lattice. A subsection of the cryo-EM map enclosing two layers of dimers in the axial direction was extracted, and tubulin dimer copies were rigid-body fitted into the subsection density. The constructed model was solvated in a triclinic water box of size 33.2 × 33.2 × 25.8 nm^3^ and subsequently neutralized with 150 mM KCl. The final MT lattice model was constructed by grafting three copies of the refined subsection models onto one another by applying a translation operation to each copy that conforms to the measured dimer periodicity.^67^

The initial model of a Dam1c subunit was derived from the cryo-EM structure of a single monomer of the Dam1c–Ndc80c complex (PDB ID: 8Q85).^34^ Missing atoms were added, and steric clashes between neighboring residues (A:511/A:516, A:499/A:503, A:517/B:296, A:374/B:216, A:447/B:283, I:55/J:59 and I:60/O:1) were resolved by choosing alternative rotamers for the affected side chains in Chimera.^68^ Short missing segments and termini within the resolved core (the Dam1 and Duo1 N-termini, the Dad2 and Spc34 internal loops, the Dad1 N- and C-termini, the Dad3, Spc34 and Ask1 N-termini, and the Hsk3 C-terminus) were completed with Modeller, retaining the cryo-EM coordinates of the Dam1 staple. The intrinsically disordered CTTs of Dam1, Duo1 and Ask1 were omitted from the initial AA model. The Dam1 and Duo1 CTTs were added at the CG model stage. Finally, side chains of the completed model were energy-minimized, and the resulting structure was used as the starting point for building the model of a Dam1c ring. The complete ring model, comprising 16 copies of Dam1c, was built by rigid-body fitting each copy into the corresponding density of the consensus map (kindly provided by Kyle Muir, University of Edinburgh, UK, and David Barford, MRC LMB Cambridge, UK). This assembled ring served as the starting model for CDMD refinement against the consensus map, following the previously published protocol.^42^

### CALVADOS simulations

The resulting AA models of the Dam1c ring and the MT were superimposed and mapped into the residue-level CALVADOS 3 representation.^49^ CG beads within folded domains were placed at the centers of mass of the corresponding residues. The unresolved Dam1, Duo1, and tubulin CTTs were generated as self-avoiding chains. Trial bead positions were rejected if they overlapped with existing beads or entered the cylindrical volume occupied by the MT. After construction of the complete CG system, the simulation box was chosen such that periodic boundary conditions along the MT axis connected the two ends of the lattice, producing an effectively infinite MT.

Consecutive beads within disordered regions were connected by harmonic bonds with an equilibrium length of 0.38 nm and a force constant of 8033 kJ mol^−1^ nm^−2^. The tertiary structure of folded domains was preserved using an elastic network model (ENM) with a force constant of 700 kJ mol^−1^ nm^−2^ between bead pairs separated by less than 0.9 nm in the reference structure. Non-bonded interactions were described by the residue-specific Ashbaugh–Hatch potential and Debye–Hü ckel electrostatics at an ionic strength of 0.15 M, unless specified otherwise. Residue protonation states were assigned for pH 7.5.

Simulations were performed using OpenMM 8.1.2^69^ in the NVT ensemble at *T* = 300 K with a Langevin integrator, a 10-fs time step, and a friction coefficient of 0.01 ps^−1^. Before production, the system was equilibrated for 5 ns with positional restraints applied to the folded regions. For umbrella-sampling simulations, all beads within the folded regions of the MT were restrained to their initial positions using harmonic restraints with a force constant of 500 kJ mol^−1^ nm^−2^. Production simulations were performed under the conditions specified for each system, with variations in ionic strength, oligomer number, and tail composition summarized in Table S1. In total, the simulations comprised approximately 4.2 ms of aggregate sampling time.

### Analysis of CALVADOS trajectories

Contacts between tubulin and the Dam1 and Duo1 CTTs were defined using a cutoff distance of *r*_*c*_ = 12 Å. For each tubulin–CTT residue pair (*i* , *j*), the contact probability *P* (*i* , *j*) was calculated as the fraction of trajectory frames in which the corresponding beads were separated by less than *r*_*c*_ (Fig. S2). To quantify the separability of the resulting contact maps, each map was approximated by its leading rank-1 component using singular value decomposition, *P* (*i* , *j*) ≈ *f* (*i*)*g* (*j*). The fraction of variance captured by the leading component was calculated as 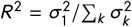. The high values obtained (*R*^2^ = 0.97–0.98, Fig. S2) indicate that the interactions are dominated by the intrinsic interaction propensities of individual tubulin and CTT residues, with little to no residue-pair specificity.

Diffusion coefficients were calculated from the displacement of the Dam1c ring along the MT axis. Global translation and rotation were removed by least-squares fitting of each trajectory frame to the initial MT structure using the folded regions of *α* - and *β* -tubulin. The center-of-mass position of each Dam1c-ring trajectory snapshot, *z* (*t*), was calculated and unwrapped across periodic boundaries. The 1D diffusion coefficient, *D* , was obtained by fitting the mean squared displacement (MSD) to MSD(*τ*) = 2*Dτ* over lag times of 10–200 ns.

To characterize CTT–tubulin contact kinetics, ten replicas were initiated from sequentially sampled frames of an equilibrated ring trajectory, with coordinates saved every 0.5 ps (Table S1). CG bead–bead distance traces for selected residues were converted into binary bound/unbound trajectories using a cutoff of *r*_*c*_ = 12 Å. Bound-state lifetimes were pooled across all 16 copies of each CTT and across all replicas and fitted to a single-exponential distribution.

The ring tilt was defined as the angle between the MT axis and the normal to a best-fit plane through the centers of mass of the 16 ring subunits.

### UCG simulations

Flaring MT ends were simulated using our previously developed UCG model.^43^ Briefly, the MT was represented by 14 coupled PFs (as in our previous work), each described as a discrete elastic rod (DER). PF stretching, bending, twisting, and twist–bending coupling were described by harmonic potentials parameterized against atomistic simulations of short GDP-tubulin oligomers.^42^ Interactions between neighboring PFs were described by Morse potentials acting between virtual lateral interaction sites (three sites per tubulin dimer). In addition, a repulsive *r*^−12^ potential was applied between the corresponding DER nodes to prevent steric overlap between neighboring PFs. The total lateral interaction potential was defined as follows:

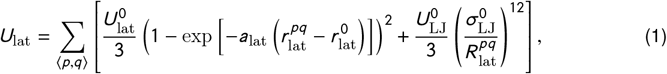

where 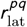 is the distance between interacting virtual sites, 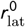 their equilibrium separation, and 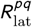 the distance between the corresponding DER nodes. The parameter *a*_lat_ controls the range of the Morse potential, while 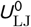 and 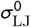 define the strength and the length scale of the repulsive term, respectively. The lateral bond strength was set to 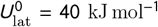, consistent with the value obtained by fitting the Morse potential to the PMF of the homotypic lateral PF–PF interaction from atomistic simulations.^43^

For the present study, our previous MT-only model^43^ was extended by a CG Dam1c ring. The ring was represented as a closed chain of *N*_ring_ = 32 spherical beads, two per Dam1c monomer, initialized as a regular polygon coaxial with the MT shaft at an initial position *z* = 5 nm relative to the lowest MT node at *z* = 0 nm. The ring bead centers were placed onto a circle of radius *R*_ring_ = *R*_MT_ + *R*_tub_ + Δ*R* ≈ 21 nm, where *R*_MT_ = 12 nm is the radius of the PF centerlines, *R*_tub_ ≈ 2.04 nm (one quarter of the 8.15-nm dimer length) is the radius of a tubulin monomer, and Δ*R* = 7 nm is the radial offset of the ring bead centers from the PF centers. The equilibrium bond length *b*_0_ = 2*R*_ring_ sin(*π*/*N*_ring_) ≈ 4.12 nm and bond angle *θ*_0_ = *π* − 2*π*/*N*_ring_ ≈ 168.75^°^ were derived from the polygon geometry, and the ring bead radius was set to *b*_0_/2 ≈ 2.06 nm so that neighboring beads are in contact, yielding inner and outer ring diameters of ∼38 nm and ∼46 nm, respectively, and a ∼5-nm offset from the MT surface, consistent with available cryo-EM structures of the Dam1c ring.^30,34^ Consecutive pairs of beads were coupled by harmonic potentials, 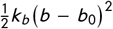, and consecutive triplets of beads were additionally coupled by harmonic angle potentials, 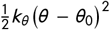, with *k*_*b*_ = 5000 kJ mol^−1^ nm^−2^ and *k*_*θ*_ = 15000 kJ mol^−1^ rad^−2^. Each ring bead interacted with every MT node through a pairwise, harmonic-repulsive potential, 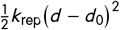 with *k*_rep_ = 500 kJ mol^−1^ nm^−2^ for center-to-center distances *d* below the contact distance *d*_0_ = *R*_tub_ + *b*_0_/2 ≈ 4.1 nm. The coordinates of ring beads were propagated with the same overdamped Brownian dynamics scheme as the MT nodes, with *D*_ring_ = 2 × 10^−5^ nm^2^ ps^−1^ per bead and Δ*t* = 10 ps.

### Time rescaling in CALVADOS simulations

CG models exhibit faster dynamics than their AA counterparts, primarily due to the reduction of degrees of freedom and the smoothing of the underlying energy landscape. By integrating out high-frequency motions and fine-grained interactions, CG potentials are effectively softer and less rugged, which lowers energy barriers and reduces frictional resistance. In addition, the absence of explicit solvent in the CALVADOS force field^47–49^ further decreases viscous drag. As a result, CALVADOS samples configurational space more rapidly, leading to an apparent acceleration of dynamical processes. A common way to assess this acceleration is to compute the acceleration factor defined as the ratio of the diffusion coefficients obtained in CG and AA simulations of the same molecular system.

Here we obtained the acceleration factor for CALVADOS as an average over three systems available in the literature^52,70,71^ (Table S2). The resulting mean acceleration factor, *a* = 127, was applied to all kinetic quantities measured in our CALVADOS simulations.

### Microtubule-end ensemble analysis and taperness parameter

Each MT end was described by the PF lengths *l*_*i*_ = *h*_*i*_ + 1, where *h*_*i*_ is the number of tubulin dimers by which PF *i* extends beyond the common base layer above the complete lattice (Fig. S4a). MT-end taperness can then be expressed as:

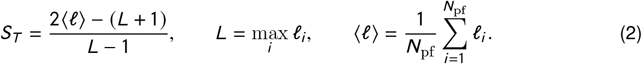

The normalized parameter *S*_*T*_ ranges from blunt to strongly tapered MT ends. For a blunt end, in which all PFs have the same length, *S*_*T*_ = 1, whereas *S*_*T*_ ≈ 0 for a uniformly tapered MT end. Negative values of *S*_*T*_ correspond to MT ends in which most PFs are substantially shorter than the longest one. The most negative value is obtained when a single PF has length *L* and all others have length 1, giving *S*_*T*_ ≈ −0.85 for a 13-PF MT lattice. Because *S*_*T*_ depends only on the mean and maximum PF lengths, it is invariant with respect to their arrangement. We used *S*_*T*_ to characterize the selected MT-end geometries. The exact structures used to calculate ⟨*F*⟩_max_ in Fig. 4b are shown in Fig. S8. These structures were generated from a fully blunt lattice by extending individual PFs with additional tubulin dimers.

To generate a representative ensemble of shortening MT ends, we sampled the length of each of the 14 PFs independently from the arc-length distribution measured in our previous cryo-ET study.^43^ The arc lengths were converted to numbers of tubulin dimers as 1 dimer ≈ 8.15 nm, and a total of 10^5^ statistically independent MT ends were generated. From this ensemble, we selected a single representative MT tip with PF heights {8, 6, 5, 4, 3, 3, 3, 2, 2, 2, 2, 1, 1, 0}, corresponding to *S*_*T*_ ≈ −0.25. For the 13-PF MT model in CALVADOS, one intermediate-length PF was removed, resulting in {8, 6, 5, 4, 3, 3, 3, 2, 2, 2, 1, 1, 0} and *S*_*T*_ = −0.23.

### Umbrella sampling

Initial configurations for umbrella sampling in CALVADOS were generated using steered molecular dynamics. For the ring–MT PMFs, the Dam1c ring was pulled along the MT axis towards the plus end, while for the Dam1c–MT PMFs, a single Dam1c monomer was pulled radially away from the MT shaft and perpendicular to the MT axis. Umbrella windows were spaced by 0.9–1.4 nm depending on the overlap between neighboring windows. In each window, the ring’s center of mass (COM) was restrained by a harmonic potential with a force constant of 10 kJ mol^−1^ nm^−2^. Lateral displacement of the ring’s COM was limited by a cylindrical flat-bottom potential with a radius of 35 nm and a force constant of 200 kJ mol^−1^ nm^−2^. Each window was simulated for 500 ns, with the ring’s COM recorded every 250 ps. To obtain the force profiles, a weighted quartic smoothing spline was fitted to the PMF using the bootstrap uncertainties as weights, and the force was calculated from its analytical derivative.

To calculate the PMFs for the dissociation of a single Dam1c monomer from the MT surface, umbrella windows were spaced by 0.39 nm. The reaction coordinate was defined as the distance in the *xy* plane between the MT’s COM and the COM of the Dam1c bridge region, comprising Dam1 residues 121–151 and Duo1 residues 150–176. A harmonic restraint with a force constant of 50 kJ mol^−1^ nm^−2^ was applied along this coordinate. Lateral displacement of the complex was limited by a cylindrical flat-bottom potential with a radius of 10 nm and a force constant of 1000 kJ mol^−1^ nm^−2^. Each umbrella window was simulated for 1 *µ*s, with the restrained coordinate recorded every 250 ps. The PMF difference was obtained by subtracting the bound-state minimum from the mean PMF in the unbound region. To convert this value to the standard binding free energy at *C* ^°^ = 1 mol L^−1^ = 1 M, the following translational entropy correction was applied:

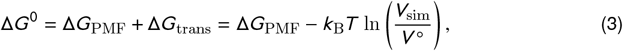

where *V*_sim_ is the volume accessible to the unbound Dam1c monomer and *V* ^°^ = 1.661 nm^3^ is the standard-state volume per molecule at 1 M.

For umbrella sampling with our UCG model, initial configurations were generated by creating replicas of the initial ring–MT configuration described above, with the ring’s COM positioned in a range from *z* = 5 nm to *z* = *z*_max_ + 16 nm, spaced by 0.5 nm, where *z*_max_ is the axial position of the highest MT node for a given PF length. In each UCG window, the ring’s COM was restrained by a harmonic potential with a force constant of 50 kJ mol^−1^ nm^−2^. Each window was simulated for 100 *µ*s, with the MT and ring coordinates recorded every 0.1 *µ*s.

All PMFs were reconstructed at *T* = 300 K using the Weighted Histogram Analysis Method^72^ implemented in GROMACS.^73^ Statistical uncertainties were estimated by bootstrapping 100 resampled profiles.

### Rupture force calculation

For each MT-tip arrangement of the representative tip, the curvature- and CTT-mediated PMFs were combined along the MT-axis reaction coordinate. Because the Dam1c CTTs can extend by up to ∼20 nm from the ring’s COM (Fig. 1c), they begin to resist ring unbinding before the ring’s COM reaches the base of the curved PFs. In contrast, curvature-mediated resistance arises only when the ring directly encounters sufficiently bent PFs. The CTT-mediated PMF was therefore shifted along the reaction coordinate relative to the curvature-mediated PMF by an offset *d* :

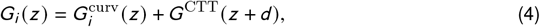

where *d* accounts for the difference in the position at which the two contributions begin to resist ring unbinding. The resulting force was insensitive to the precise value of this offset: varying *d* between 5 and 15 nm (Fig. S7) changed the ensemble force by less than 1 pN.

To compute the one-second rupture force, *F*_1*s*_ , each combined PMF was first modified by the applied force as *G*_*i*_ (*z*) – *F z* . The force-dependent activation barrier, 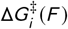, was defined as the largest free-energy ascent along the tilted profile. The corresponding escape rate was then calculated as:

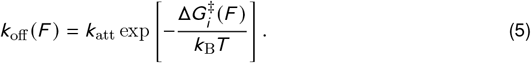

The attempt rate was approximated by the diffusive relaxation rate of the bound well,

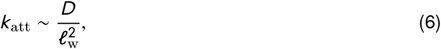

where *D* = 0.01 *µ*m^2^ s^−1^ is the experimental ring diffusion coefficient along the MT lattice,^12^ and 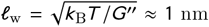 is the characteristic width of the bound well, with *G"* denoting the average curvature of the PMF at its minimum.

## Contributions

M.K. and M.I. conceptualized the project. M.K. performed the simulations, analyzed the data, and drafted the manuscript. M.I. and H.G. secured funding and supervised the project. All authors reviewed and edited the final manuscript.

## Acknowledgments

All authors acknowledge the support provided by the Max Planck Society. M.K., M.I., and H.G. additionally thank the Deutsche Forschungsgemeinschaft (DFG): Project-ID 449750155 – RTG 2756, Project A1 for supporting this research. M.I. acknowledges the funding provided by the Royal Society (University Research Fellowship, URF*\*R1*\*251325) and the Academy of Medical Sciences (Springboard Award, SBF0010*\*1119). We thank Kyle Muir (University of Edinburgh, UK) and David Barford (MRC LMB Cambridge, UK) for sharing the cryo-EM data that enabled the construction of the initial models and helpful clarifications. We also thank Carter J. Wilson (MPI-NAT, Germany) for valuable and insightful discussions.

## Code availability

Initial AA/CG coordinates, CALVADOS setup configurations, analysis scripts, PMF profiles, the extended custom DER code including the Dam1c ring, as well as the raw CALVADOS and UCG trajectories will be available from the corresponding author upon reasonable request. All remaining data and figures supporting the findings of our study are included as Supporting Information (Figs. S1–S8, Tables S1–S2, Movie S1).

