## Supporting Information for "Dynamic microtubule-end structure governs multivalent kinetochore coupling by the Dam1c ring"

Movie S1: Visualization of the Dam1c ring diffusion along the MT lattice. Color coding as in Fig. 1a.

Table S1: Summary of the CALVADOS simulations.

| Oligomer size $N$ | CTTs | Oligomers per replica | Ionic strength (M) | Replicas | Avg. time per replica | Frame interval |
|---|---|---|---|---|---|---|
| 1–3 | All | 4 | 0.15 | 3 | ~5 $\mu$s | 1 ns |
| 4 | All | 3 | 0.15 | 4 | ~5 $\mu$s | 1 ns |
| 5–7 | All | 2 | 0.15 | 5 | ~5 $\mu$s | 1 ns |
| 8–15 | All | 1 | 0.15 | 10 | ~5 $\mu$s | 1 ns |
| 16 | All | 1 | 0.01 | 40 | ~10 $\mu$s | 1 ns |
| 16 | All | 1 | 0.05 | 40 | ~10 $\mu$s | 1 ns |
| 16 | All | 1 | 0.10 | 40 | ~10 $\mu$s | 1 ns |
| 16 | All | 1 | 0.20 | 40 | ~10 $\mu$s | 1 ns |
| 16 | All | 1 | 0.25 | 40 | ~10 $\mu$s | 1 ns |
| 16 | All | 1 | 0.15 | 40 | ~10 $\mu$s | 1 ns |
| 16 | $\Delta$Tub | 1 | 0.15 | 40 | ~10 $\mu$s | 1 ns |
| 16 | $\Delta$Dam1 | 1 | 0.15 | 40 | ~10 $\mu$s | 1 ns |
| 16 | $\Delta$Duo1 | 1 | 0.15 | 40 | ~10 $\mu$s | 1 ns |
| 16 | $\Delta$Dam1 + $\Delta$Duo1 | 1 | 0.15 | 40 | ~10 $\mu$s | 1 ns |
| 16 | $\Delta$All | 1 | 0.15 | 40 | ~10 $\mu$s | 1 ns |
| 16 | All | 1 | 0.15 | 10 | 200 ns | 0.5 ps |

Table S2: Overview of literature systems and their extracted diffusion coefficients used to quantify dynamical scaling between AA and CG representations.

| System | $D_{\mathrm{AA}}$, $\mathrm{nm}^2/\mathrm{ns}$ | $D_{\mathrm{CG}}$, $\mathrm{nm}^2/\mathrm{ns}$ | Acceleration $a$ |
|---|---|---|---|
| ProT$\alpha$ – H1 dimer[52] | 0.071 ± 0.003 | 8.32 ± 0.48 | 117 |
| Lge1 single chain[70] | 0.12 | 15.47 ± 0.87 | 128 |
| Hst5 (A-Disp)[71] | 0.16 ± 0.02 | 21.38 ± 0.76 | 136 |

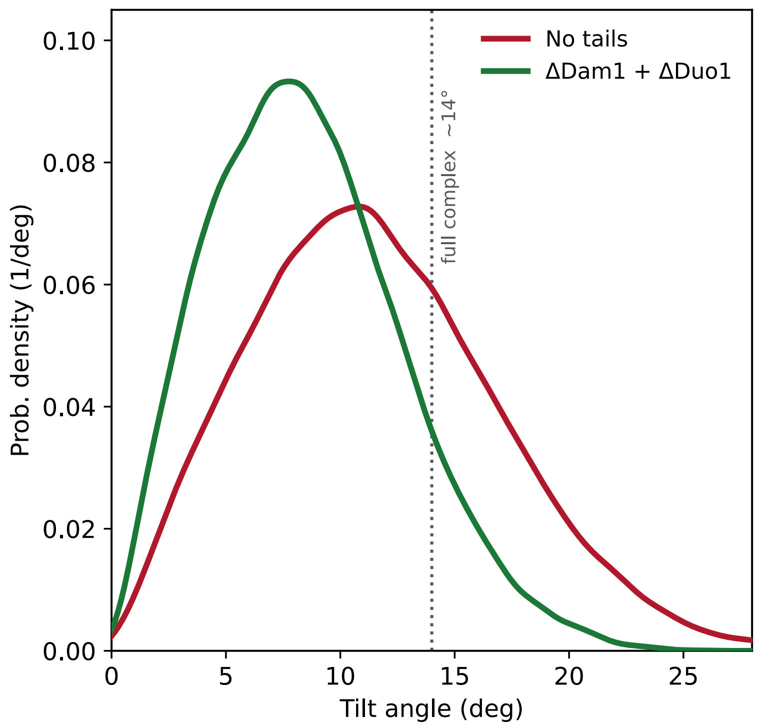


Figure S1: **Probability densities of the tilt angle of the Dam1c ring relative to the MT axis for two CTT-deletion variants.** A ring with all CTTs removed (no tails) versus a ring after removing Dam1 and Duo1 CTTs (ΔDam1 + ΔDuo1). The dotted vertical line marks the mean tilt of the full complex.

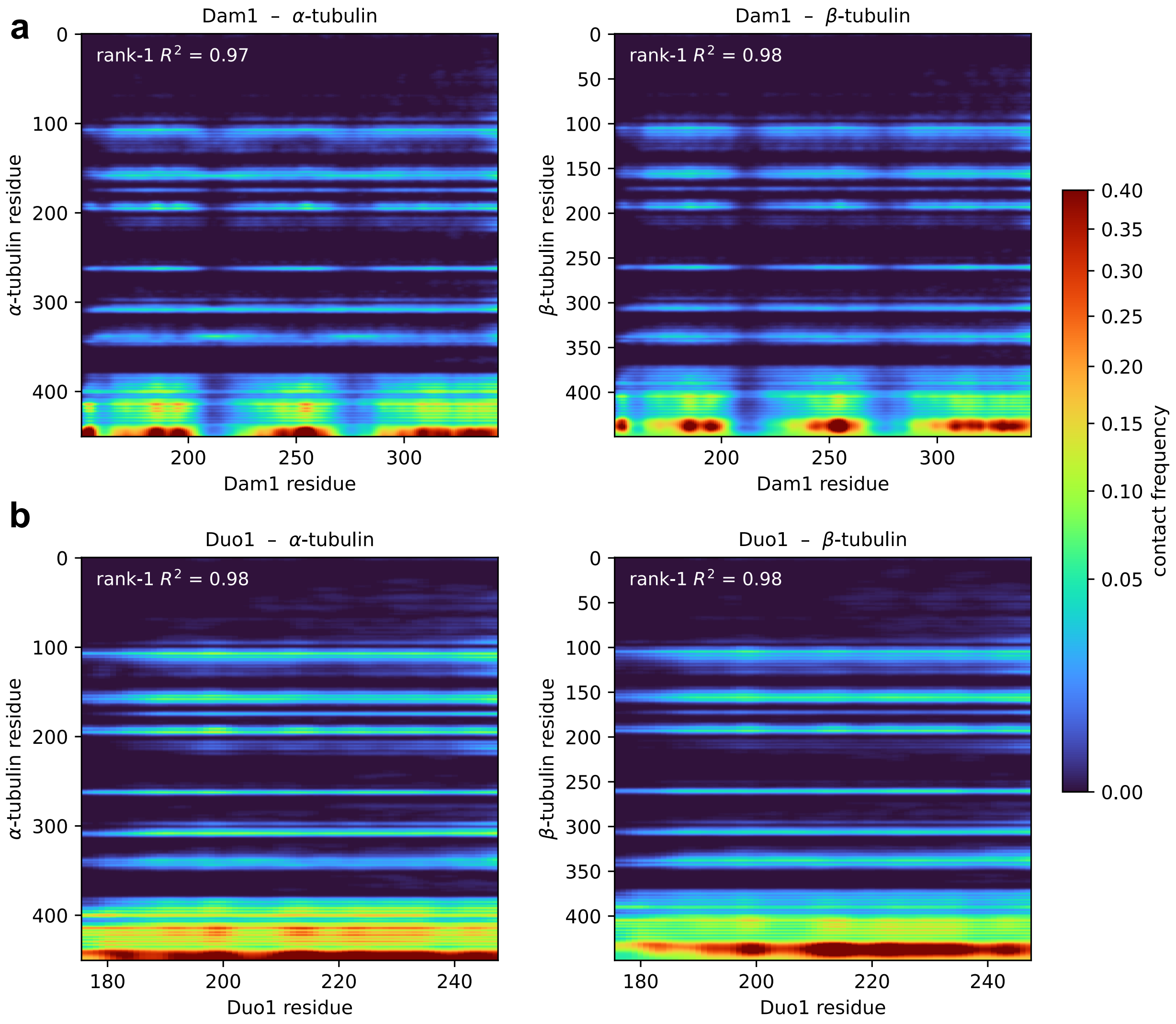


Figure S2: **Two-dimensional residue–residue contact probability maps $P(i, j)$ between the Dam1c CTTs and tubulin. (a)** Dam1 CTT with $\alpha$-tubulin (left) and $\beta$-tubulin (right). **(b)** Duo1 CTT with $\alpha$-tubulin (left) and $\beta$-tubulin (right). The color scale gives the contact frequency. Insets report the fraction of variance $R^2$ captured by the leading rank-1 component of each map, obtained by singular value decomposition.

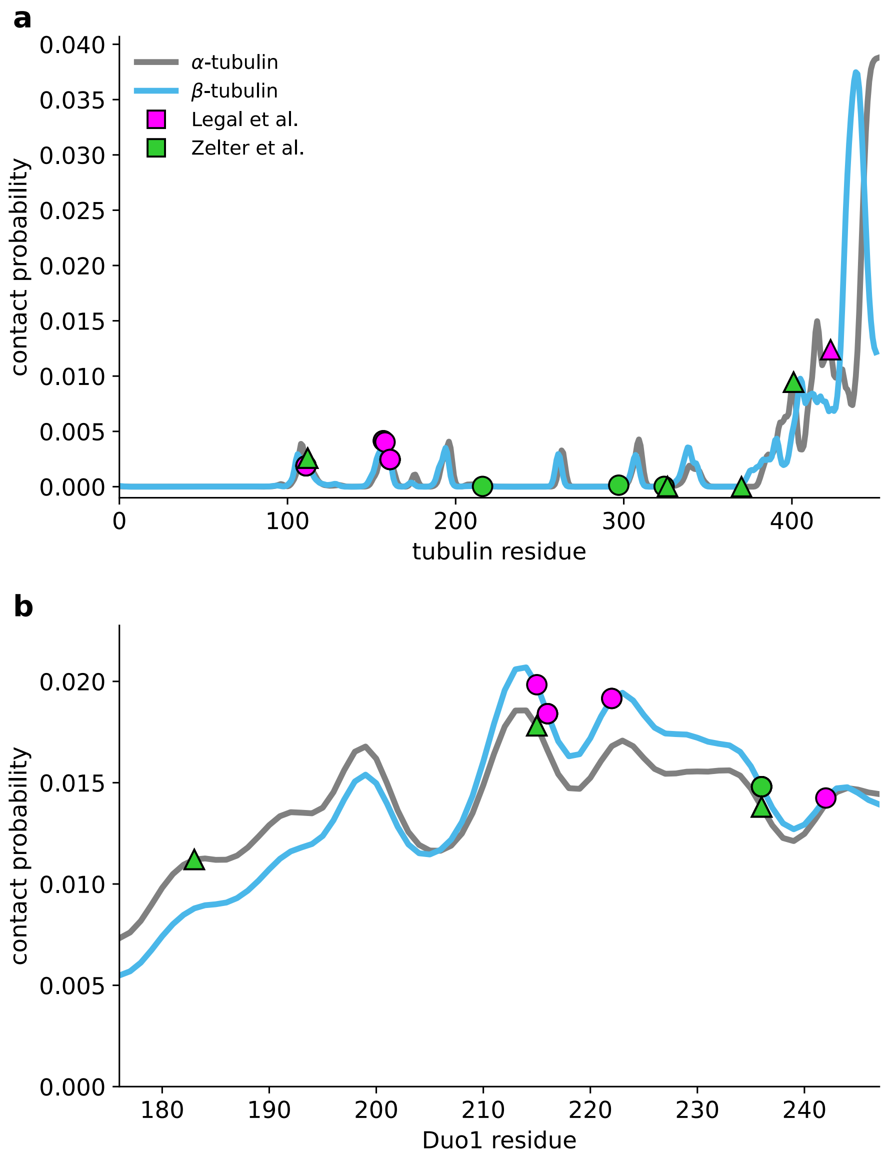


Figure S3: **Marginal contact probabilities for the Duo1 CTTs. (a)** Per-residue marginal contact probability along $\alpha$-tubulin (gray) and $\beta$-tubulin (blue). **(b)** Per-residue marginal contact probability along the Duo1 CTT. Residues identified in cross-linking assays by Legal *et al.* (K–E/D, magenta)[29] and Zelter *et al.* (K–K, green)[28] are shown with symbols.

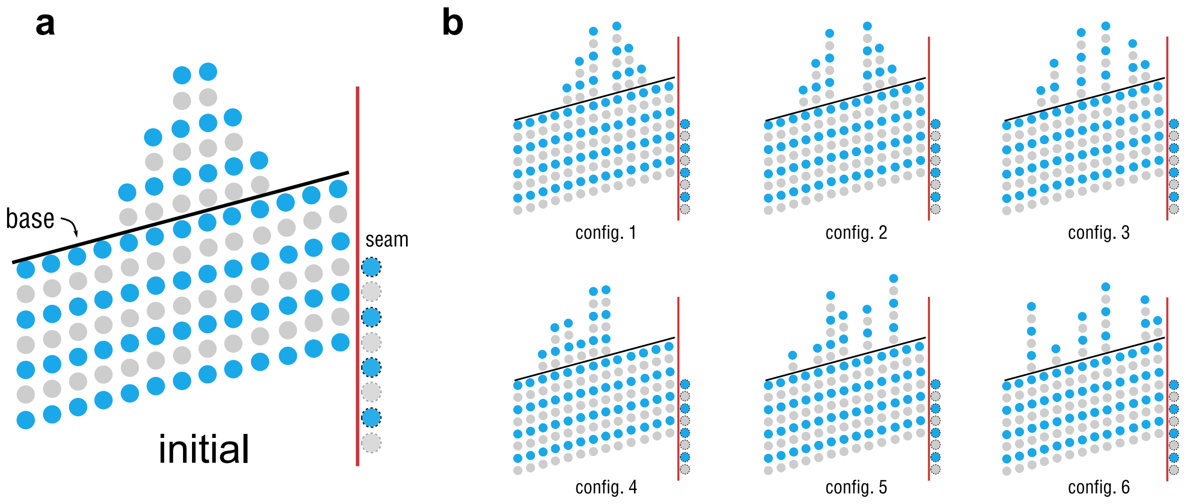


Figure S4: **Unrolled schematics of the 13-PF MT ends.** The tubulin monomers of each PF are shown as blue and gray circles. The black line marks the common base of the complete lattice from which the PF heights $h_i$ are counted, and the red line marking the lattice seam. **(a)** The initial MT-end structure, illustrating the representation used throughout. **(b)** The six permutations of the same set of PF lengths (configurations 1–6). The corresponding CTT-mediated force profiles are shown in Fig. S5.

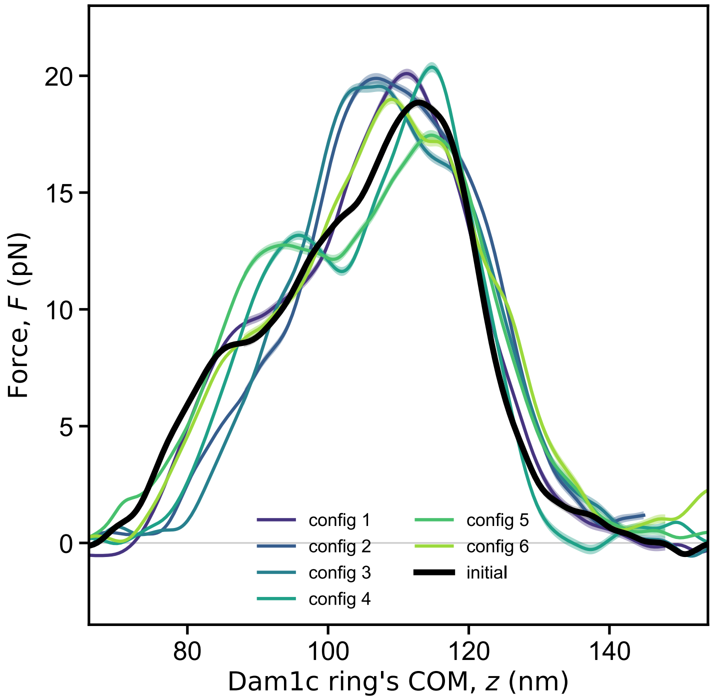


Figure S5: **Mean resistance force acting on the Dam1c ring as a function of the position of its center-of-mass along the MT axis.** Colored curves correspond to six PF permutations (configurations 1–6) of the same distribution of PF lengths. The black curve is the initial arrangement. Shaded bands represent statistical uncertainty. See Fig. S4 for the exact compositions.

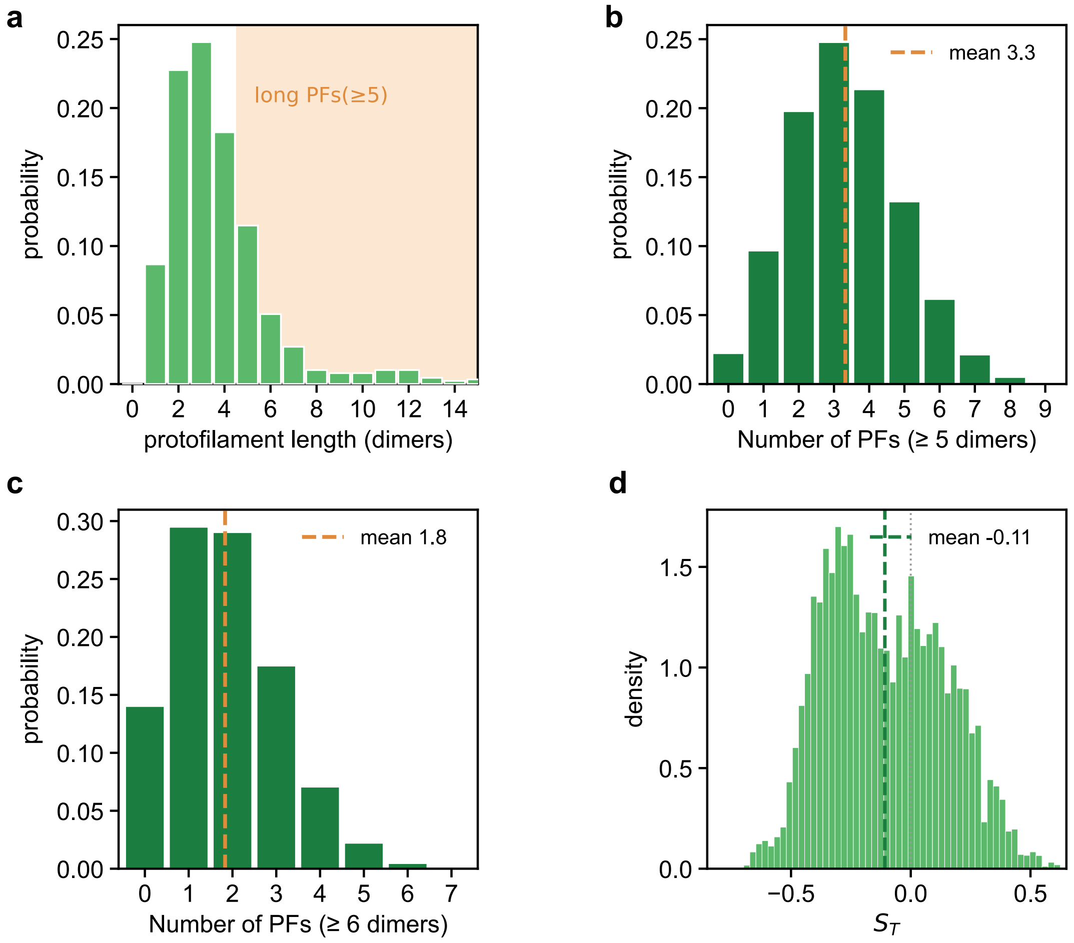


Figure S6: **Ensemble of MT-end geometries sampled from cryo-ET. (a)** Distribution of individual PF lengths obtained from Kalutskii *et al.*[43] **(b)** Distribution of the number of long PFs ($\geq 5$ dimers) per MT end. **(c)** Same for PFs of $\geq 6$ dimers. **(d)** Distribution of the MT-end shape parameter $S_T$ (Eq. 2).

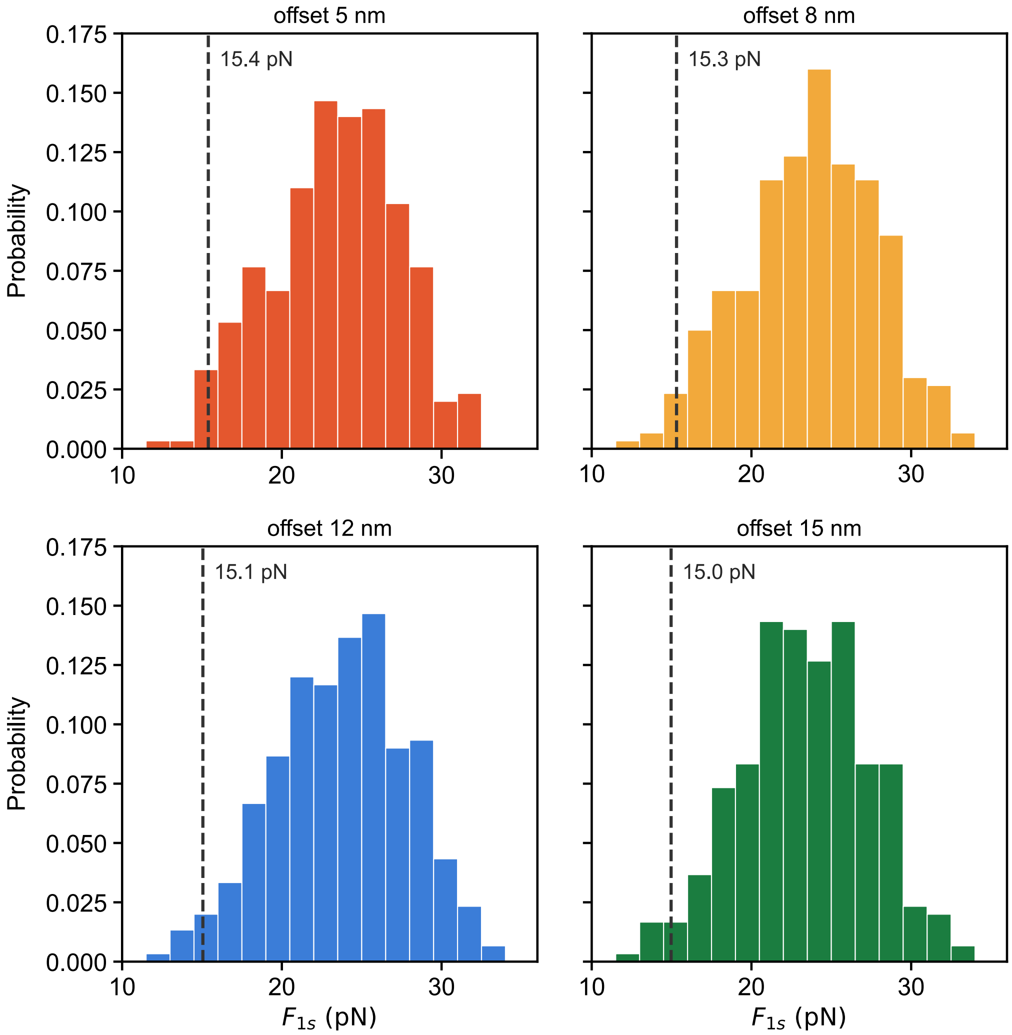


Figure S7: **Distributions of the one-second rupture force $F_{1s}$.** Each panel corresponds to a different offset $d$ used when combining the CTT- and curvature-mediated PMFs (5, 8, 12 and 15 nm). The dashed line in each panel marks the ensemble-averaged rupture force.

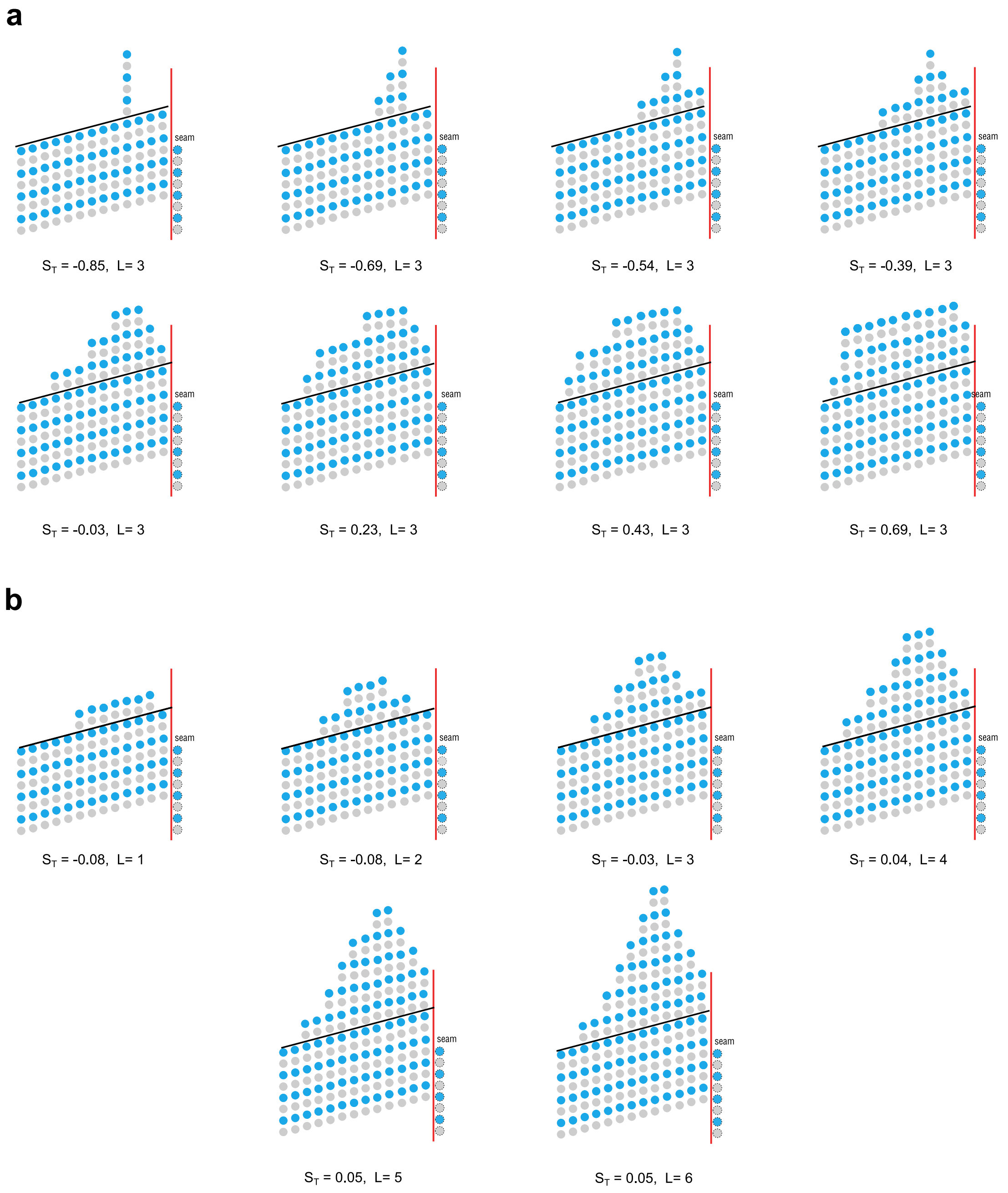


Figure S8: **Composition of the MT-end structures probed in our CALVADOS and UCG simulations. (a)** The eight tip geometries used for the $S_T$ scan in Fig. 4b. The PF taper length was fixed at $L_T = 3$ dimers, while $S_T$ was varied from $-0.85$ to 0.69. The fully blunt tip, in which all PFs have a length of three dimers ($S_T = 1$), is not shown. **(b)** The six tip geometries used for the taper-length scan of Fig. 4b. Representation as in Fig. S4.
